# Automatic cell type harmonization and integration across Human Cell Atlas datasets

**DOI:** 10.1101/2023.05.01.538994

**Authors:** Chuan Xu, Martin Prete, Simone Webb, Laura Jardine, Benjamin J. Stewart, Regina Hoo, Peng He, Kerstin Meyer, Sarah A. Teichmann

## Abstract

Harmonizing cell types across the single-cell community and assembling them into a common framework is central to building a standardized Human Cell Atlas. Here we present CellHint, a predictive clustering tree-based tool to resolve cell type differences in annotation resolution and technical biases across datasets. CellHint accurately quantifies cell-cell transcriptomic similarities and places cell types into a relationship graph that hierarchically defines shared and unique cell subtypes. Application to multiple immune datasets recapitulates expert-curated annotations. CellHint also reveals underexplored relationships between healthy and diseased lung cell states in eight diseases. Furthermore, we present a workflow for fast cross-dataset integration guided by harmonized cell types and cell hierarchy, which uncovers underappreciated cell types in adult human hippocampus. Finally, we apply CellHint to 12 tissues from 38 datasets, providing a deeply curated cross-tissue database with ∼3.7 million cells and various machine learning models for automatic cell annotation across human tissues.

## Introduction

The past decade has seen an accumulation of single-cell genomics datasets profiling a variety of tissues at different developmental stages. By integrating these resources, international consortia such as the Human Cell Atlas (HCA) have begun to chart a standardized reference map for the human body.^1, 2^ A challenge in assembling distinct cell atlas datasets is that different laboratories use their own definitions of cell types, often resulting in inconsistencies in the naming schema. This challenge is being partially met by the ongoing Cell Ontology database effort, which aims at unifying cell type labels across communities.^3^ Other contributions come from the NIH Human BioMolecular Atlas Program (HuBMAP)^4^ project and its ASCT+B tables that name and interlink major anatomical structures, cell types and biomarkers,^5^ and the Cell Annotation Platform (a collaborative effort funded by Schmidt Futures, https://celltype.info). However, standardizing cell types across datasets, denoted here as the process of cell type harmonization, has intrinsic challenges beyond nomenclature. These include the variability in cell quality caused by various technical confounders, presence of shared versus novel cell types across datasets, and the discrepancy in annotation resolution due to differences in cell sampling depth and in laboratory-specific annotation strategies.

A number of label transfer methods have been developed to match annotated cell types across datasets.^6–13^ A limitation of many of these methods is that they rely strongly on the pairwise dataset alignment under a reference-to-query scheme. Aligning multiple datasets simultaneously is still arduous as it requires multiple pairwise comparisons and brings challenges in information integration and consistent visualization. A further limitation lies in the nature of machine learning-based models that, without a prebuilt cell type hierarchy, neglect the relationships among cell types both within and across datasets and usually lead to “flat” classifiers.

Furthermore, current efforts for single-cell data integration largely focus on correcting batch effects across datasets.^9, 14–18^ Information relating to cell types annotated by individual studies is not fully utilized in batch effect regression owing to naming inconsistencies. Once cell types are successfully harmonized across datasets, data integration can be supervised in an annotation-aware manner. Although deep learning-based methods such as scANVI^19^ and scGen^20^ are able to include cell type information in the integration pipeline, further efforts are needed to control for the degree of intervention from cell annotations, and to incorporate cell type relationships into the data integration process.

To meet these challenges, we develop an automated workflow for cell type harmonization connected with cross-dataset integration. Specifically, we develop a predictive clustering tree (PCT)-based tool, CellHint, to efficiently align multiple datasets by assessing their cell-cell similarities and harmonizing cell annotations. Based on this, CellHint defines semantic relationships among cell types and captures their underlying biological hierarchies, which are further leveraged to guide the downstream data integration at different levels of annotation granularity. We have applied this pipeline to 49 datasets, confirmed its effectiveness in data harmonization and integration, and provided a collection of organ atlases and machine learning models to the wider community for automatic cell type annotation.

## Results

### An automated workflow for cell type harmonization and integration

CellHint infers cell type relationships across datasets in two main steps: predicting the distances among cells and summarizing the alignments among cell types (**Table S1; STAR Methods**). CellHint first calculates the transcriptional distances between cells and cell types within a single reference dataset, and then builds a clustering tree for this dataset using PCT, a multi-target regression tree algorithm.^21^ Different from classical multi-output decision trees, with PCT the dependency among cell types in the reference dataset is considered. To reduce runtime and avoid overfitting, during the top-down induction of PCT, the reference tree is pruned at the cell taxonomy (i.e., node) where further splitting will result in two homogeneous cell states based on an F-test. Next, each query cell from the other datasets will be assigned to the matched tree leaf according to its expression pattern, and based on this, the dissimilarities between this cell and cell types in the reference dataset are predicted to be the prototype of the corresponding leaf which usually contains a cluster of transcriptionally similar cells (**Figure 1**). In short, once a PCT is built for the reference dataset, this information is used to predict a query cell’s dissimilarity profile across reference cell types, thereby effectively scaling the query datasets to the reference. By iteratively building such a tree using each dataset as a reference and mapping the other datasets onto it, CellHint derives a global distance matrix representing the inferred dissimilarities between all cells and cell types (i.e., calculated or predicted transcriptional distances within or across datasets, respectively). Since this algorithm has no assumption about gene expression distribution and cell-cell distances are predicted within the span of the clustering trees, CellHint is able to produce batch-insensitive dissimilarity measures, enabling a robust cross-dataset meta-analysis.

**Figure 1.**
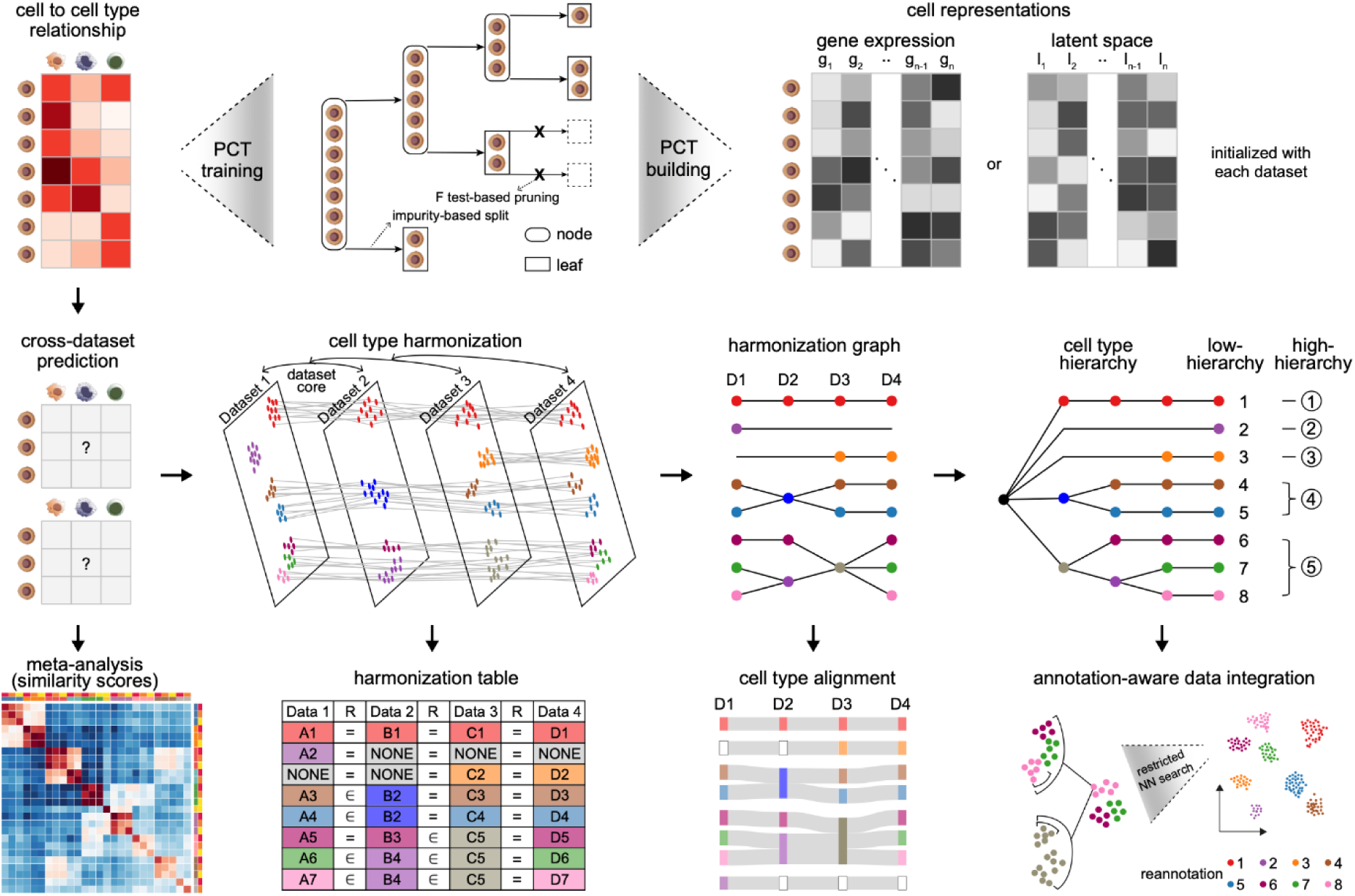
Framework of CellHint. For each dataset, CellHint calculates a cell-to-cell-type dissimilarity matrix based on the gene expression space or low-dimensional latent space. Next, CellHint builds a predictive clustering tree, based on which the transcriptomic dissimilarities between cell types in the reference and cells from the remaining datasets can be predicted. By iteratively considering each dataset as the reference, CellHint derives a global cell-to-cell-type dissimilarity matrix and enables two downstream analyses, including a batch-insensitive meta-analysis and construction of a harmonization graph. The latter is achieved by cell type alignments between two starting datasets, followed by iterative addition of new datasets and extension of cell type relations, leading to a semantically defined harmonization table. This harmonization graph can be reorganized to obtain two levels of cell type hierarchies, which serve as input for the annotation-aware data integration to restrict the neighborhood search space. PCT, predictive clustering tree; NN, nearest neighbors. See also Table S1.

Next, the resulting dissimilarities between cells and cell types are integrated into a harmonization graph by considering all possible pairs of reference-to-query predictions (**Figure 1**). Specifically, we define a dataset core which summarizes original cell types into semantically connected ones including matched (“is equivalent to” or “=”) and divisible (“contains” or “∋”, and “belongs to” or “∈” for one-to-multiple and multiple-to-one alignments, respectively) cell types. Through iterative addition of a new dataset, this core is gradually enlarged to re-define an expanded cell type repertoire, namely, the CellHint harmonization table (**Figure 1; STAR Methods**). The advantage of this procedure is that the order of datasets added has a minimal impact on the structure of the harmonization graph. As a result, cell types in the graph can be reorganized to approximate the underlying biological hierarchies including the high and low hierarchies (**Figure 1**). This also represents an avenue to integrate knowledge across studies by connecting their manual and biologically meaningful groupings. Moreover, to improve sensitivity in discovering dataset-specific cell types, CellHint defines two levels of novelties for cell types in a given dataset: unmatched cell types (“NONE”) which cannot align with any cell type from the other datasets, and unharmonized cell types (“UNRESOLVED”) which fail to integrate into the harmonization graph after the final iteration.

CellHint further utilizes the cell type relationships to supervise the single-cell data integration (**Figure 1**). Given the harmonized cell types across datasets, the neighborhood graph is built by searching the neighbors of each cell in a restricted manner within the same cell type group which is determined by combining the cell type hierarchy with the inferred transcriptomic dissimilarities (**STAR Methods**). By controlling the size of cell type groups, CellHint can adjust the extent of supervision from cell annotations. Data scalability and efficiency are also anticipated to be improved as the neighborhood search space is dramatically narrowed down.

### CellHint harmonization recapitulates expert-curated annotations

To validate the harmonization pipeline in CellHint, we selected five immune datasets which were initially included in the CellTypist database and deeply curated by experts into consistent cell types.^8^ These datasets had varying cell type compositions, developmental stages, tissue coverages, and transcriptome sparsities, presenting a real-world scenario for cell type harmonization (**Figure S1A**). As expected, we observed substantial batch effects when visualizing the 417,866 cells onto the uniform manifold approximation and projection (UMAP)^22^ (**Figure S1B**). We thus asked whether CellHint can harmonize the cell types of the datasets to recapitulate the ground-truth annotations (**Figure S1C**).

Despite the various confounding factors present in the five datasets, CellHint was able to reconstruct their cell type relationships from two complementary angles. Firstly, CellHint predicted the transcriptomic distances among the 129 original cell types, and based on this, revealed a high degree of cross-dataset cell type replicability (**Figure S1D**). Specifically, unsupervised hierarchical clustering using the inferred distances not only grouped the same cell types from different datasets regardless of their batch confounders, but also demonstrated a biological structure comprising compartments of hematopoietic progenitors, myeloid, and lymphoid cells. This meta-analysis also uncovered finer structures underlying these datasets such as the innate lymphoid cells and T cells, the latter of which were further subdivided into CD4+ αβ, CD8+ αβ, and γδ T cells (**Figure S1D**). Secondly, CellHint harmonized the 129 cell types into 66 at the low-hierarchy level, identifying both dataset-shared and -specific cell types (**Figures 2A and S1E**). Reorganization of the harmonization graph further categorized them into 52 high-hierarchy cell groups (**Figures 2B and S1F**).

**Figure 2.**
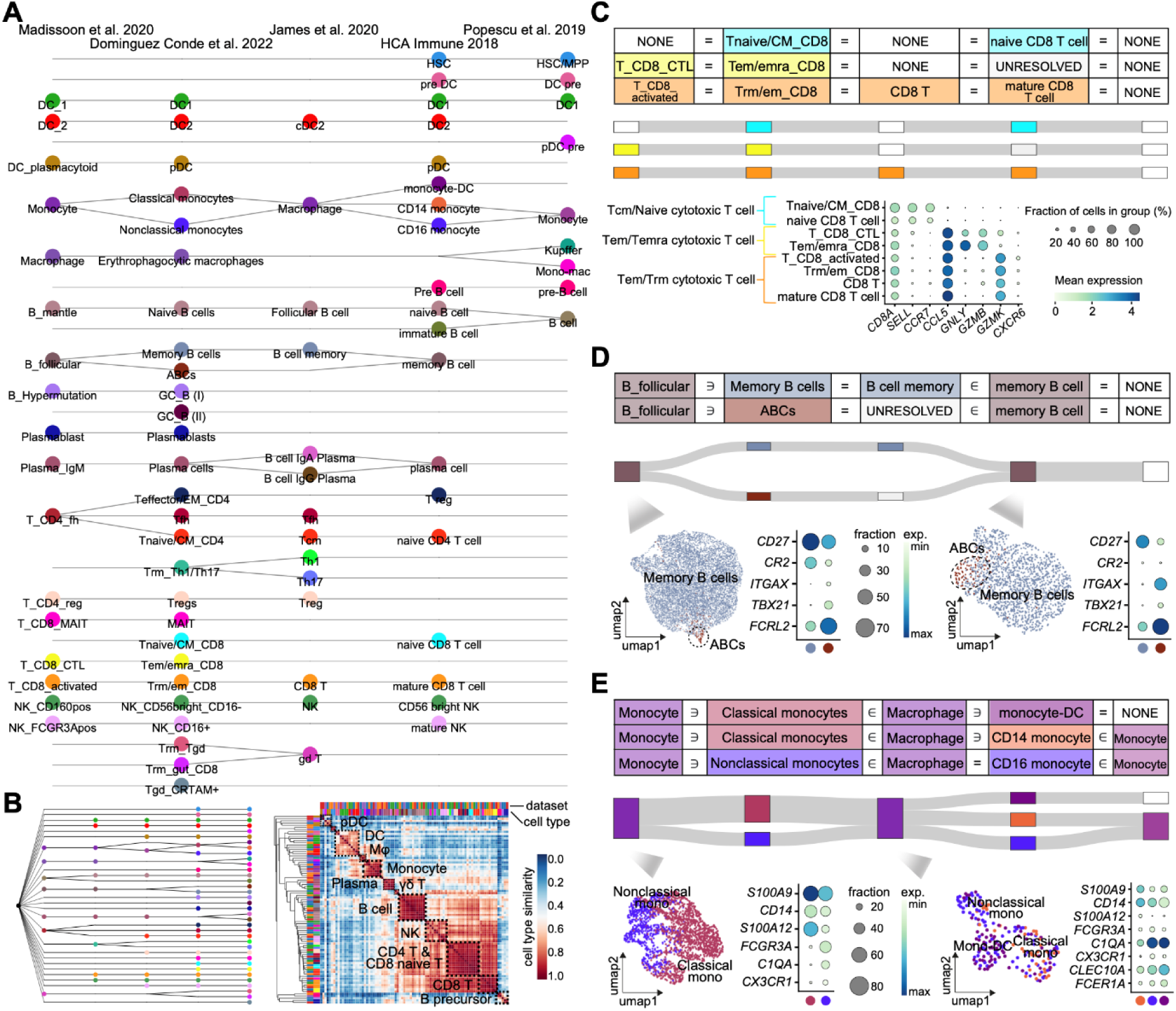
CellHint harmonization recapitulates manual annotations across five immune datasets. (A) Harmonization graph showing selected cell type relationships across five immune datasets shown on top. Cell type labels are colored according to their low-hierarchy alignments. Note that the lines in the graph reflect the connections among transcriptionally similar cell types regardless of their names. The order of datasets applies to (C), (D), and (E). (B) Left: cell type hierarchy reorganized from (A). Right: heat map displaying the unsupervised hierarchical clustering of cell types based on their inferred similarities by CellHint. Cell types are colored as in (A), and clustered into cell compartments marked in the plot. (C) Upper: part of the harmonization table containing eight CD8+ T cell populations, as well as their alignments visualized as a Sankey diagram. Lower: dot plot showing expression of marker genes across the eight cell populations corresponding to three subtypes harmonized by CellHint. Color of the dot represents normalized gene expression, and size represents the percentage of cells expressing a given gene. (D) Upper: as in (C), but for the five memory B cell-related populations. Lower: UMAP visualizations of memory B cells in two datasets that contain a hidden population of age-associated B cells (ABCs). Dot plots show expression of marker genes for memory B cells and ABCs, with the color and size representing normalized gene expression and the percentage of cells expressing a given gene, respectively. (E) Upper: as in (D), but for the eight monocyte-related cell types. Lower: as in (D), but for the two cell populations consisting of classical and nonclassical monocytes (left), and of three monocyte subtypes (right), respectively. See also Figures S1, S2, and S3.

As well as the harmonizations that could be confirmed at the nomenclature level (e.g., “DC_2”, “DC2”, and “cDC2”), CellHint resolved complex cell type alignments not obvious based on their nomenclatures. For example, CellHint harmonized the eight annotations of CD8+ αβ T cells into three cell states (**Figure 2C**). These included a reassuring match between “Tnaive/CM_CD8” and “naive CD8 T cell”, as well as two matches not easily resolvable using nomenclature only: between “T_CD8_CTL” and “Tem/emra_CD8”; and among “T_CD8_activated”, “Trm/em_CD8”, “CD8 T”, and “mature CD8 T cell”. Examination of marker gene expression validated these cell identities aligned (**Figure 2C**). Similarly, for natural killer (NK) cells, CellHint harmonized seven NK annotations into two subtypes as supported by their respective gene signatures, that is, the CD16+ (“NK_FCGR3Apos”, “NK_CD16+”, and “mature NK”) and CD16- (“NK_CD160pos”, “NK_CD56bright_CD16-”, “NK”, and “CD56 bright NK”) NK cells (**Figure S2A**).

A number of cell types were hierarchically harmonized, leading to the identification of cell subtypes not annotated as such in the original publications (**Figure 2A**). In one of the datasets, we previously described a specialized and rare memory B cell population, age-associated B cells (ABCs), and extrapolated its presence to other datasets by means of manual inspection.^8^ Using the automated harmonization algorithm in CellHint, we reproduced these findings by corroborating the existence of ABCs in two other datasets profiling the adult spleen and bone marrow, respectively^23^ (**Figure 2D**). The other example was the “B cell” population identified in the fetal liver,^24^ which we demonstrated to consist of a spectrum of transitional subpopulations from immature to naive B cells (**Figure S2B**). Moreover, the harmonization module in CellHint is able to detect homogeneous subtypes which may not readily be separated using clustering-based approaches. In line with our manually curated results, we found three transcriptionally similar myeloid populations (“monocyte-DC”, “CD14 monocyte”, and “CD16 monocyte”) collectively forming the “Macrophage” annotated in the gut dataset^25^ (**Figure 2E**), as well as three distinct CD4+ T cell populations (“Tnaive/CM_CD4”, “Tfh”, and “Teffector/EM_CD4”) constituting a single heterogeneous “T_CD4_fh” cell type in a spleen dataset^23^ (**Figure S2C**).

Interestingly, CellHint discovered new cell types that were previously overlooked. For instance, the γδ T cells (“gd T”) initially annotated in the gut dataset^25^ were revealed by CellHint to contain a hidden population of epithelium-resident CD8+ αβ T cells (“Trm_gut_CD8”) which closely resembled the γδ T cells (**Figure S2D**). This was also the case for plasma cells defined in two datasets^8, 23^ where CellHint hierarchically divided them into the IgA (“B cell IgA Plasma”) and IgG (“B cell IgG Plasma”) plasma subpopulations based on another dataset^25^ (**Figure S2E**).

To further test the stability of our algorithm in multi-dataset alignment, we permuted the order of datasets incorporated during harmonization and confirmed the robustness of the aforementioned observations (**Figures S2G and S2H**). Taken together, through comparisons with expert-graded annotations across multiple immune datasets, we showcase the capacity of CellHint to perform meta-analysis, harmonize cell types, define cell hierarchies, and inform potential new biology.

### Application of CellHint to other omics datasets

We next tested the capability of CellHint to deal with datasets beyond the context of single-cell transcriptomics. To this end, we collected five single-cell and single-nucleus datasets^26–28^ generated using the assay for transposase-accessible chromatin with sequencing (scATAC-seq and snATAC-seq) for profiling the blood, bone marrow, lymph node and lung (**Figure S3A**). To compare cell types and hierarchies with those derived from transcriptomics (**Figures 2A and S1E**), we focused on the 119,046 immune and progenitor cells corresponding to 94 original cell annotations from these chromatin accessibility datasets (**Figures S3A and S3B**).

Using CellHint, we revealed 42 and 55 cell types at the high- and low-hierarchy levels, respectively, with the vast majority of them echoing the transcriptome-based cell type alignments (**Figures S3C and S3D**). Examples are the cross-dataset correspondence for cell types such as dendritic, B, and T cells. This analysis also located sub-branches of cell types not found in the transcriptome-based cell hierarchy. For example, the hematopoietic stem and progenitor cells (“HSPC”) were harmonized by CellHint into two categories: the megakaryocyte/erythrocyte progenitors (“MK/E prog”) and hematopoietic stem cells (“HSC”), with the former further subdivided into megakaryocyte-erythroid progenitors (“MEP”) and erythroid cells (“Ery”) and the latter split into “HSC” and hematopoietic multipotent progenitors (“HMP”) (**Figure S3D**). We also observed a misalignment between “CD4 TEM” and “ILC3/Th17”, which may result from a potential decrease in the ability of scATAC-seq datasets to define robust cell identities due to frequent peak dropouts, which is less of an issue in transcriptomic datasets.^29, 30^

### CellHint disentangles disease-enriched cell states

Cell diversity is often increased in pathological conditions due to the emergence of novel cell types or molecular reorganization of the same cell types.^31, 32^ To test the utility of CellHint in these contexts, we applied it to four single-cell datasets^33–36^ profiling the adult human lung, collectively covering 479,135 cells from 130 individuals in health plus eight diseases resulting in pulmonary fibrosis: idiopathic pulmonary fibrosis (IPF), chronic obstructive pulmonary disease (COPD), nonspecific interstitial pneumonia (NSIP), chronic hypersensitivity pneumonitis (cHP), interstitial lung disease (ILD), sarcoidosis (SA), systemic sclerosis (SSc), and polymyositis (PM) (**Figure S4A**).

Prominent technical batch effects existed across these datasets, making the discovery of disease patterns challenging (**Figures S4B and S4C**). Despite this, the harmonization graph allowed us to locate cell types and subtypes missed by individual studies and to interpret pathological cell states by extending their contexts across studies. We observed that the aberrant basaloid cells (“Aberrant_Basaloid”), a pathogenic population lining the myofibroblast foci in the IPF lung,^33^ were aligned with the distal alveolar epithelial population (“KRT5-/KRT17+”) enriched in pulmonary fibrosis^35^ (**Figure 3A**), suggestive of a single aberrant population named differently between studies. This was further corroborated by an independent analysis using logistic regression-based label transfer. In addition, the harmonization graph indicated mixing of this population into the “AT1” cell type of a third dataset.^36^ *In silico*, we were able to resolve these populations and verified their presence by combinatorial expression of multiple marker genes (**Figure 3A**). Similarly, goblet cells in the diseased state from two of the datasets^33, 34^ were demonstrated to be heterogeneous, divisible into *MUC5B*+ and *SCGB3A2*+*SCGB1A1*+ populations (**Figure 3B**). CellHint can further refine the original cell annotations to match their underlying biology. For instance, the IPF “Fibroblast” in Adams et al.^33^ was shown to align with the lipofibroblast-like subpopulation, as supported by its overexpression of *HAS1* and *PLIN2* (**Figure S4D**). The IPF “Fibroblast” in Morse et al.,^34^ on the other hand, was *ACTA2*+ and corresponded to the activated myofibroblasts during pulmonary fibrosis (**Figure 3C**). Other examples included the macrophage (**Figure 3D**) and T cell (**Figure S4E**) subpopulations harmonized by CellHint and confirmed at the molecular level. All these highlight the power of CellHint in enhancing the specificity and interpretability of cell types across diseases and studies.

**Figure 3.**
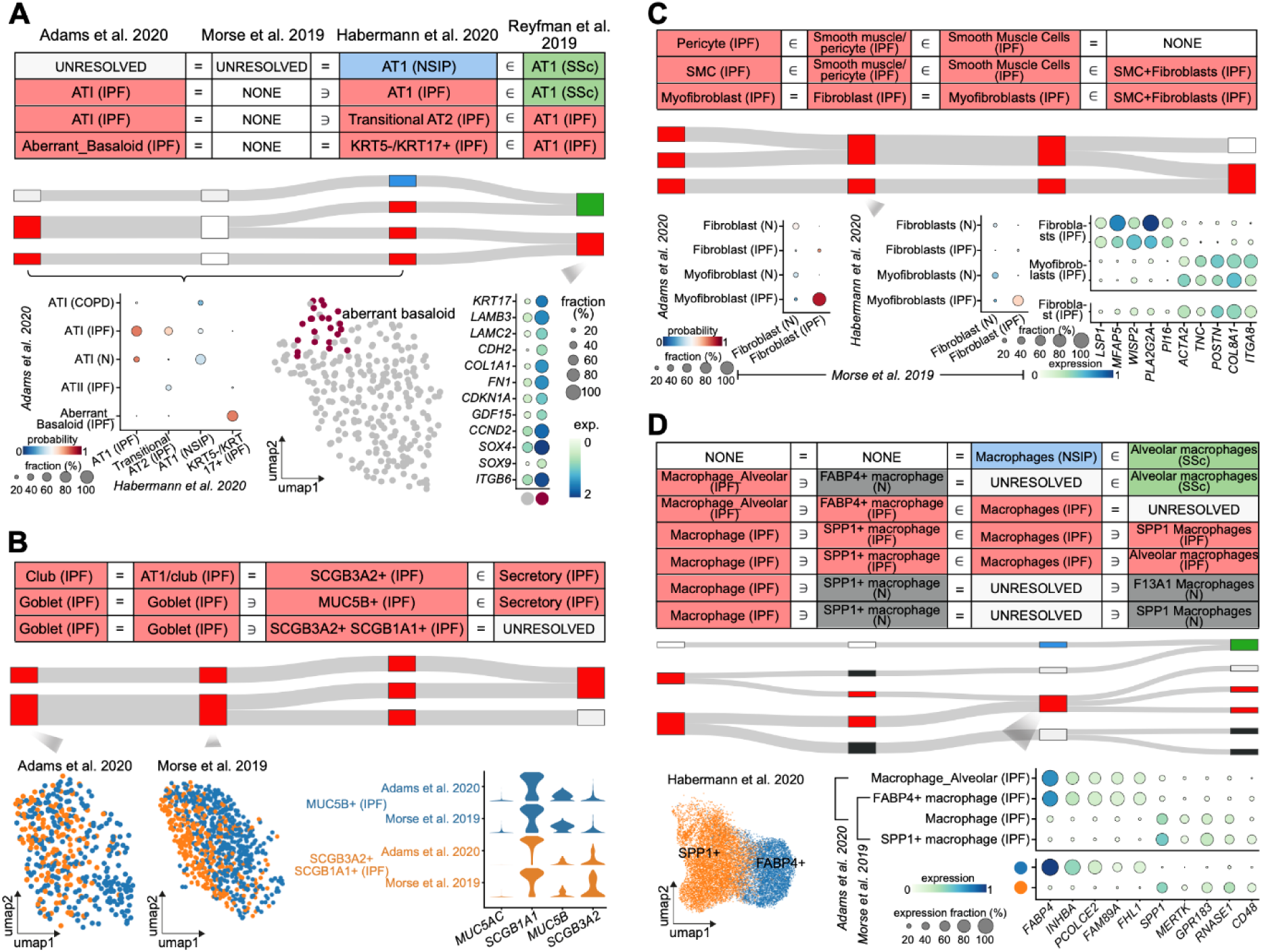
Examples of pathological cell states harmonized by CellHint. (A) Upper: part of the harmonization table containing alveolar epithelial populations, as well as their alignments visualized as a Sankey diagram. Lower: scatter plot showing cell type label transfer from the first dataset to the third dataset, highlighting a correspondence between “Aberrant_Basaloid” and “KRT5-/KRT17+”. Color and size of the dot denote the mean prediction probability and cell type alignment fraction, respectively. Dot plot to the right shows expression of marker genes of the aberrant basaloid cells separated from the “AT1” cells of the fourth dataset, with color and size of the dot representing normalized gene expression and percentage of cells expressing a given gene, respectively. The order of datasets shown on top applies to (B), (C), and (D). (B) Upper: as in (A), but for the pathological secretory cells. Lower: UMAP plots showing goblet cells in two datasets that are harmonized into two subtypes based on the third dataset. Violin plot to the right displays marker gene expression across the two goblet subtypes. (C) Upper: as in (A), but for the collagen-producing cell types. Lower: as in (A), but for the IPF “Fibroblast” in the second dataset that is harmonized into myofibroblasts based on the first and third datasets. Dot plot to the right shows expression of marker genes for the “Fibroblast” identified (bottom) as compared to those from the other two datasets (top). (D) Upper: as in (A), but for macrophage-related populations in the four datasets. Lower: as in (B), but for the pathological “Macrophages” in the third dataset that are harmonized into two macrophage subtypes. Dot plot to the right shows expression of marker genes for the two subtypes identified (bottom) as compared to those from the other two datasets (top). See also Figure S4.

With CellHint, epithelial and stromal cell types in the four datasets were consistently harmonized, in a manner that discriminated the transcriptomic differences not only among cell subtypes but also among disease statuses (**Figures 4A, 4B**, **S4F, and S4G**). For example, the basal, ciliated, and alveolar epithelial cell types were clearly separated into healthy and diseased cell states, with most of the disease statuses further distinguishable in the harmonization graph. Conversely, in the immune cell compartment, many cell types were harmonized irrespective of their disease contexts, such as the partial alignment between normal and COPD mast cells, as well as among NK cells in normal, COPD, NSIP and IPF conditions (**Figures S4H and S4I**). These observations were in line with the clinical manifestations of pulmonary fibrosis by which epithelial and stromal cells were most affected,^37^ while the relatively weaker transcriptomic changes in immune cell types were insufficient to drive a cell identity transition.

**Figure 4.**
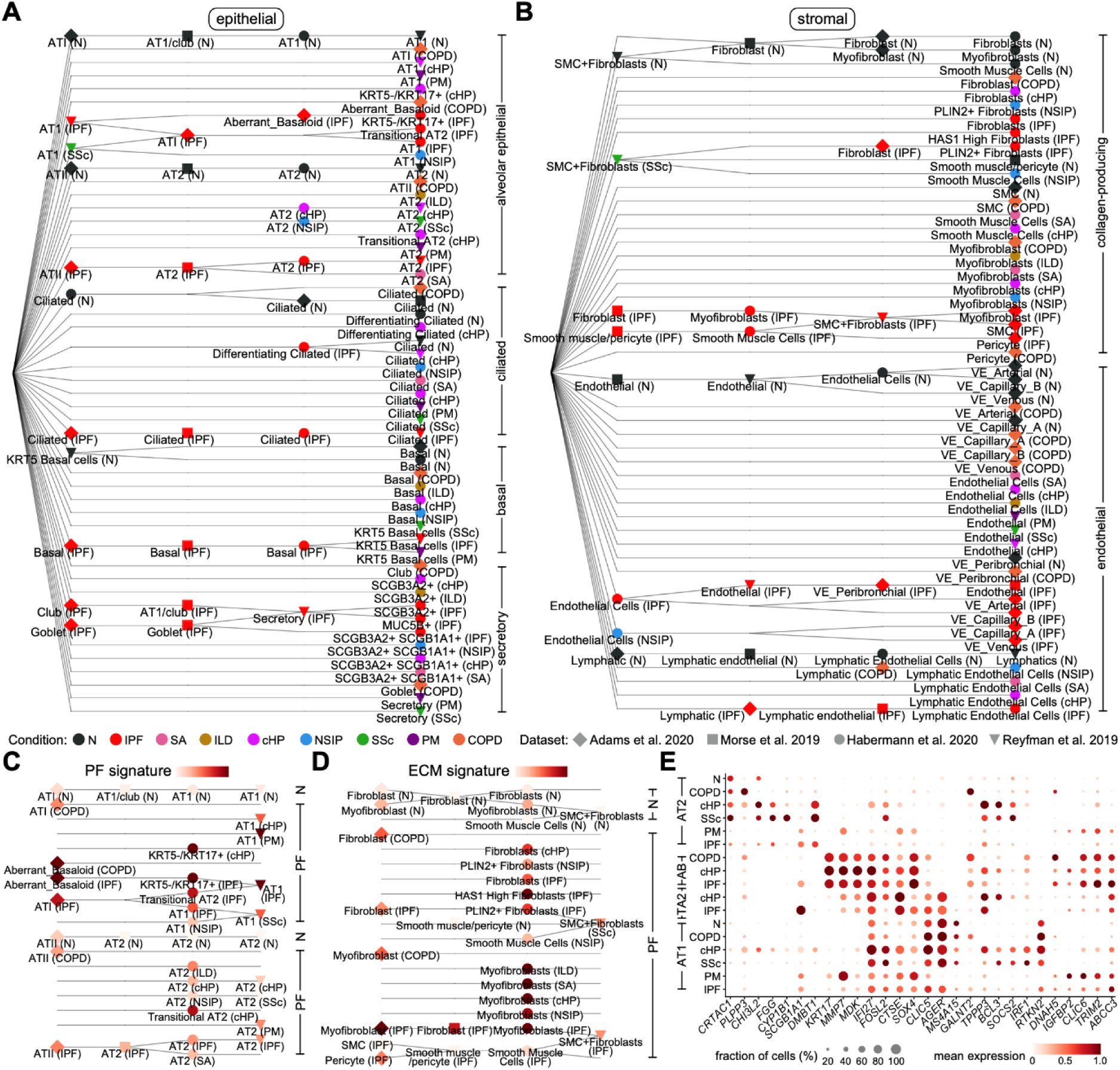
CellHint harmonizes diseased cell states for discovery of molecular alterations. (A) Cell type hierarchy visualizing the harmonized epithelial cell types across the four lung datasets. Color and shape of the dot represent the disease condition and dataset of origin, respectively. (B) As in (A), but for harmonized stromal cell types across the four lung datasets. (C) Distribution of the pulmonary fibrosis (PF)-defining gene signature in the harmonization graph focusing on the alveolar epithelial cell types. Color and shape of the dot represent the degree of PF signature and dataset of origin, respectively. (D) As in (C), but for distribution of the extracellular matrix (ECM) signature in collagen-producing cell types. (E) Dot plot showing expression of differentially expressed genes among alveolar epithelial cell types and diseases, with the color and size representing normalized gene expression and the percentage of cells expressing a given gene, respectively. Cell types and diseases are grouped across datasets based on their alignments inferred by CellHint. AB, aberrant basaloid cells; TA2, transitional AT2 cells. See also Figure S5.

Furthermore, harmonization of all cell types into one unified representation contributes to the goal of investigating cellular pathodevelopmental trajectories. We used a list of pulmonary fibrosis-defining biomarkers^33, 35^ as signature genes to quantify the degree of epithelial remodeling in the lung. Among alveolar epithelial cell types, this signature was particularly prominent in the aberrant basaloid cells, as expected. In addition to this, the alveolar type 1 (AT1) and 2 (AT2) cells were scored higher in fibrotic states of the eight diseases as compared to the healthy controls, potentially accounting for their disease-specific alignments (**Figure 4C**). Moreover, consistent with the CellHint alignment where a proportion of transitional AT2 cells were intermixed within the AT1 cells in the IPF lung, these two cell types exhibited higher fibrosis scores than healthy and diseased AT2 cells. This indicated an impaired alveolar epithelial regeneration process from AT2, with transitional AT2 cells developing into AT1 cells^38^ (**Figure 4C**).

For airway epithelial cells, overrepresentation of the pulmonary fibrosis signature was observed in the secretory cell compartment. These included diseased club cells that aligned across datasets, and IPF-related goblet cells, in particular the *MUC5B*+ subtype harmonized in three of the datasets (**Figure S5A**). Among the collagen-producing stromal cells, an excessive accumulation of extracellular matrix (ECM) was detected in pathological fibroblasts and myofibroblasts (**Figure 4D**). Between them, myofibroblasts in five diseases (ILD, SA, cHP, NSIP, and IPF) had activation signatures, and showed a much stronger enrichment of ECM genes than healthy and diseased fibroblasts, including the IPF “Fibroblast” in Morse et al.^34^ which was harmonized into the myofibroblast population (**Figure 3C**). In the lung endothelium, we located a match between the peribronchial endothelial cells from Adams et al.,^33^ and all IPF endothelial cells in the other three datasets that were *COL15A1*+, a marker suggestive of a widespread bronchiolization occurring in endothelial cells of the distal lung in IPF (**Figure S5B**). Lastly, we assessed the distribution of the profibrotic gene signature^33^ across harmonized immune cells, and found its enrichment in the *SPP1*+ macrophages, and to a lesser extent, the *F13A1*+ macrophages, in comparison to the *FCN1*+ macrophages (**Figure S5C**). This was consistent with the reported role of this *SPP1*+ macrophage population in activating myofibroblasts during fibrosis.^39^

We next grouped cell types and disease conditions according to their alignments in the cell type hierarchy. Through this, duplicated information can be consolidated and molecular signals can be systematically interrogated. In alveolar epithelial cells, the AT2-transitional AT2-AT1 molecular axis was maintained across the various disease states; at the same time, specific disease variations in this trajectory were detected (**Figure 4E**). We found activation of genes such as *BCL3* and *SOCS2* in AT1 and AT2 cells from cHP and SSc, as compared to cells from PM and IPF. Interestingly, *BCL3* has been reported to alleviate pulmonary injury in both mice and humans,^40, 41^ and the observed downregulation in the most severe form of pulmonary fibrosis (i.e., IPF) may reflect its association with disease severity. In contrast, compared to other diseases, PM and IPF induced overexpression of several other genes in AT1 and AT2 cells, including *IGFBP2*, a biomarker of IPF,^42^ *CLIC6*, a key gene upregulated in IPF,^43^ and *TRIM2*, part of the microRNA-369-3p/TRIM2 axis relevant for regulating pulmonary fibrosis.^44^

Transcriptional alterations were also detectable in a more cell type- and disease-restricted manner (**Figure 4E**). For instance, in AT2 cells, expression of *PLPP3* and *GALNT2* was largely specific to cells in COPD, whereas *CYP1B1* was detected as uniquely expressed in SSc, a chronic autoimmune disease affecting multiple organs including the kidney and lung (and inhibition of *CYP1B1* has been shown to prevent fibrosis in the kidney^45^). We also observed a partial loss of the AT1 cell identity specifically in PM, illustrated by the deficiency in expression of AT1 markers *AGER* and *RTKN2*, as well as a gain of expression in known pulmonary fibrosis marker *MMP7*.^46^ Further extending this analysis to the secretory cells (**Figure S5D**) and fibroblasts (**Figure S5E**), we characterized both disease-shared and -specific alterations in transcriptomes of multiple cell types and subtypes.

In summary, application of CellHint to four diseased lung datasets successfully harmonized the cell types in different compartments, disentangled the aberrant cell states in multiple diseases, revealed their underlying pathological signatures, and facilitated a comprehensive investigation of transcriptomic changes.

### Cell re-annotation and data integration by CellHint

Given the harmonized cell types across datasets, CellHint can further refine the identity of each cell by reannotating it within the context of multiple datasets. Specifically, on the basis of cell harmonization information, including cell identities annotated in the dataset of origin and inferred from the other datasets, each cell can be reannotated to one of the harmonized cell types in the harmonization graph corresponding to a given row in the harmonization table (**Figure 1**). CellHint reannotated the cells from the five immune datasets into 66 low-hierarchy and 52 high-hierarchy cell types, providing precise classifications with different granularities for individual cells (**Figure 5A**). For instance, the previously annotated γδ T cells were grouped as a single cell type at the high hierarchy and were further classified into two low-hierarchy cell types: 88% as γδ T cells (“NONE = Trm_Tgd ∈ gd T = NONE = NONE”) and 12% as epithelium-resident CD8+ T cells (“NONE = Trm_gut_CD8 ∈ gd T = NONE = NONE”).

**Figure 5.**
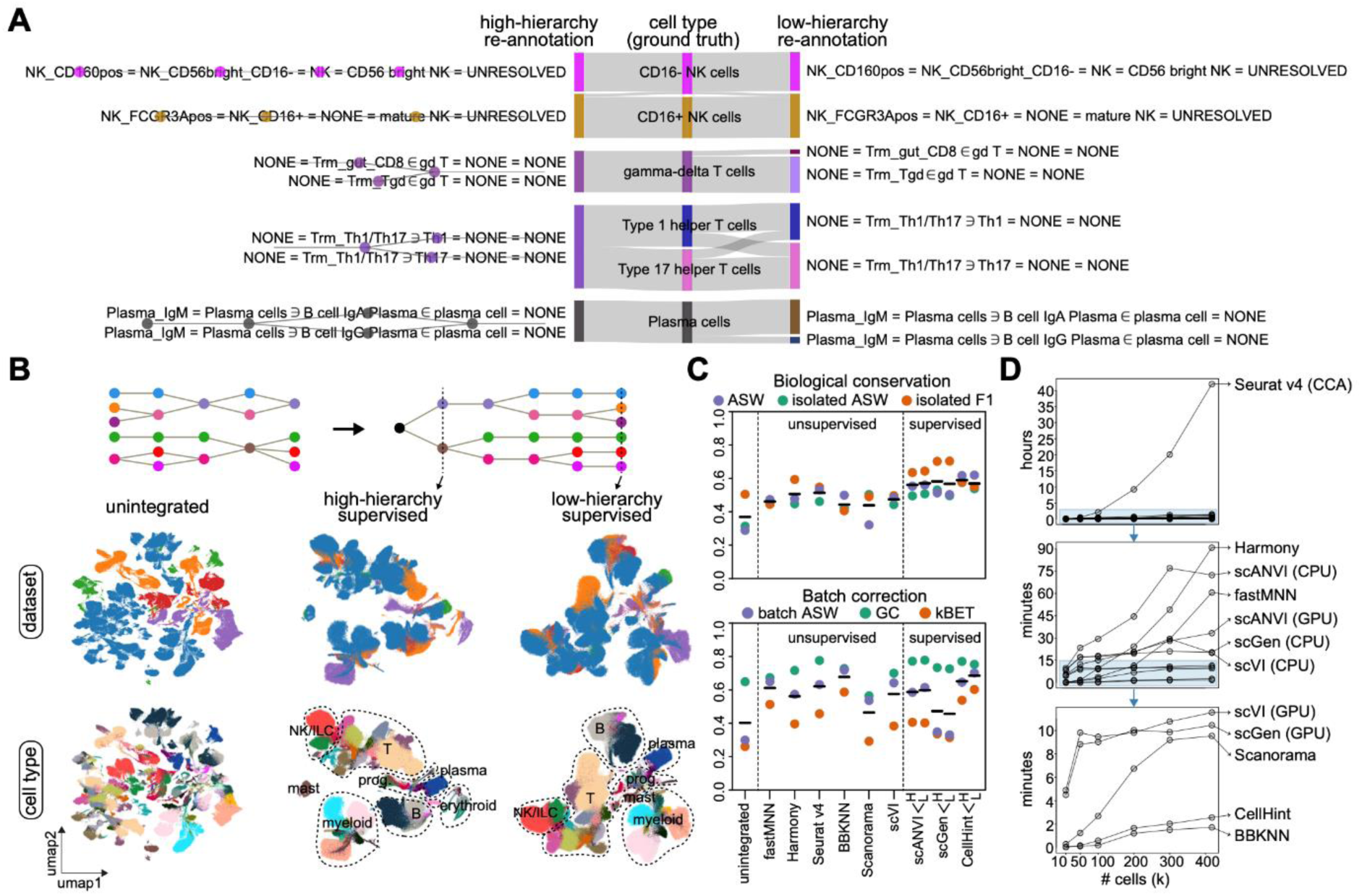
CellHint enables cell reannotation and supervised data integration. (A) Example Sankey diagram depicting the correspondence among the high-hierarchy cell reannotation (left), the ground-truth annotation (middle), and the low-hierarchy cell reannotation (right) from five immune datasets. Cells are reannotated by concatenating original cell type labels across datasets, corresponding to rows designated by the harmonization table. (B) Upper: schematic of the harmonization graph and cell type hierarchy used as input for CellHint-based data integration. Lower: UMAP representations of cells from the five immune datasets overlaid by the dataset of origin (top) and cell type (bottom). Transcriptomic structures are shown for unintegrated cells (left), and cells integrated with supervision from high-hierarchy (middle) and low-hierarchy (right) cell types. Color codes of datasets and cell types can be found in Figure S6A. (C) Assessment of biological conservation (upper) and batch removal (lower) after data integration using the six unsupervised and three supervised methods shown on bottom. For supervised methods, both high- and low-hierarchy annotations are used. Horizontal black lines denote the averages of the integration metrics. ASW, average silhouette width; GC, graph connectivity; kBET, k-nearest-neighbor batch-effect test. (D) Comparisons of runtime (y axis) across the nine integration methods, based on cell subsets ranging from 10,000 to 400,000 (x axis). For data integration using scVI, scANVI and scGen, both CPU and GPU are tested. See also Figures S6 and S7.

Through reannotation, all cells were placed into a common naming schema, allowing data integration to be supervised by annotations. In particular, the cell hierarchies built under this schema, in combination with the transcriptional distances, offered an accurate and efficient approach to construct the neighborhood graph. This confined neighbor search to cells from the same branches or similar cell types, thereby purifying cell neighbors at a minimal search cost. Notably, the extent to which the cell hierarchies guide data integration can be adjusted through controlling the number of neighboring cell types (i.e., meta neighbors) that are incorporated into the search space. This approach also applies to partially annotated datasets where local transcriptomic structures can be tuned using annotated cells only. We employed this strategy to integrate cells from the five immune datasets, and based on this, revealed the expected transcriptomic organization of immune cells (**Figures 5B and S6A**).

To benchmark CellHint against widely used data integration methods, we compared the performance of CellHint on the five immune datasets with that of other methods, including six unsupervised (FastMNN, Harmony, Seurat v4, BBKNN, Scanorama, and scVI) and two supervised (scANVI and scGen) ones^9, 14–20^ (**Figure S6B**). A set of integration metrics were used to assess the performance of these tools (**STAR Methods**). With respect to the biological conservation, annotation-aware methods (scANVI, scGen, and CellHint) showed better preservation of cell types across datasets than unsupervised ones, regardless of use of low- or high-hierarchy annotations (**Figure 5C**). The degrees of batch removal, on the other hand, were comparable across most methods, with the top performers being Seurat v4, BBKNN, and CellHint. The performance of CellHint was further confirmed by benchmarking on the four diseased lung datasets (**Figure S7**).

**Figure 7.**
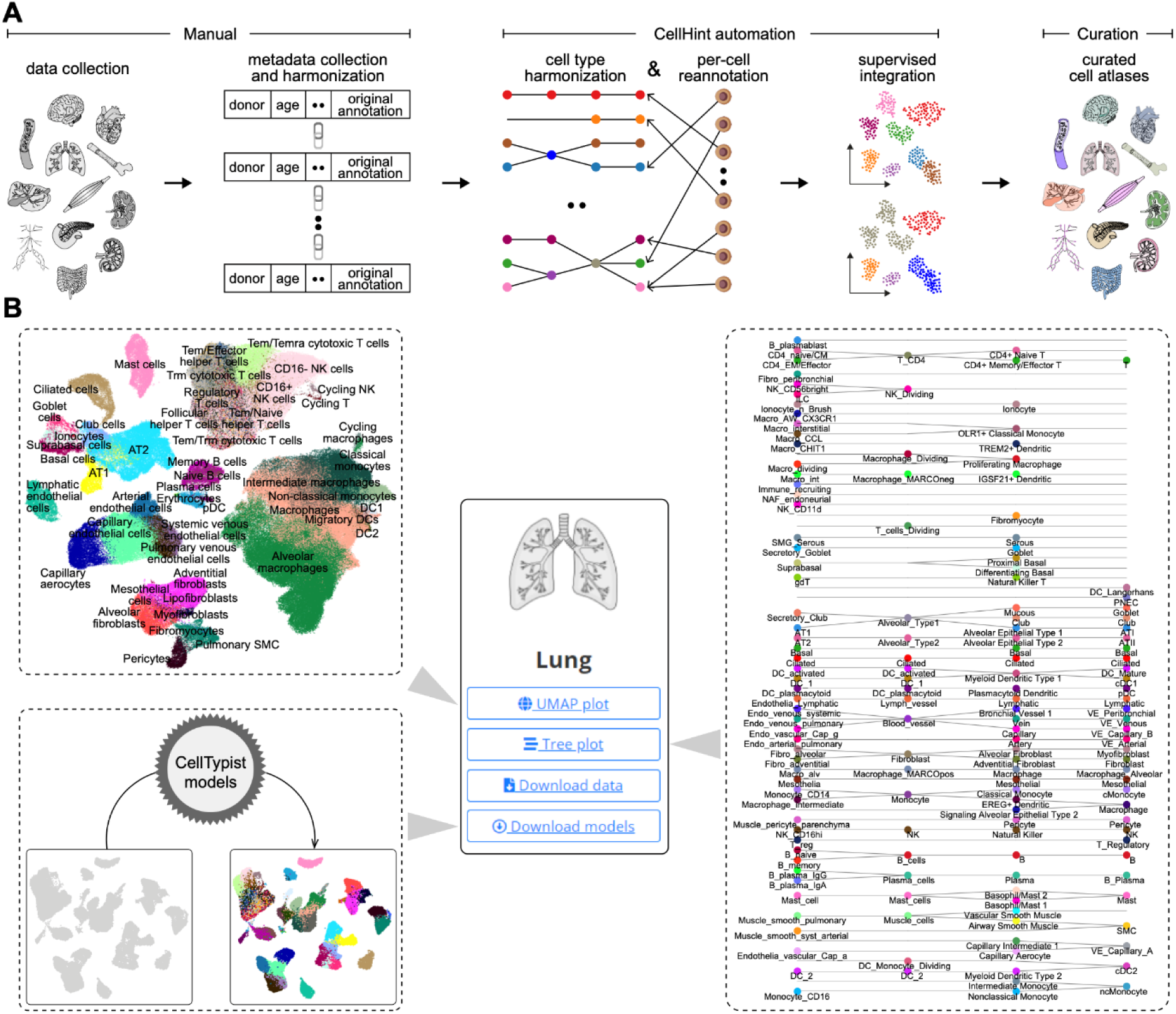
A standardized multi-organ resource of cell type harmonization, integration, and classification. (A) Workflow of building standardized organ atlases using CellHint, from collection of data and metadata, cell type harmonization, cell reannotation, supervised data integration, to the final manual curations. (B) Illustration of the three modules in our multi-organ resource, including integrated atlases with harmonized and curated cell types (top left), harmonization graphs with cross-dataset relationships among cell types (right), and light-weight machine learning models for automated cell classifications (bottom left). See also Table S2.

Since only part of the dataset is exploited when anchoring neighbors to query cells, we also expect an improved efficiency of data integration by using CellHint. To test this, we benchmarked the runtime of all nine methods based on different subsets of the five immune datasets generated by downsampling cells from 10,000 to 400,000 (**Figure 5D**). Five methods were able to integrate these cells within 15 minutes: scVI and scGen (using graphics processing unit [GPU]), Scanorama, CellHint, and BBKNN. Among them, CellHint and BBKNN were the only two methods that integrated more than 400,000 cells in two minutes.

In conclusion, after reannotating cells through the CellHint harmonization module, CellHint further provides an integration module to tune the data structure towards harmonized cell types whilst mitigating the effects from batch confounders in an efficient manner.

### CellHint assembles an integrated atlas of adult human hippocampus

To further demonstrate the biological insights that can be offered by cell type harmonization and integration, we used CellHint to assemble six single-nucleus transcriptomics datasets^47–52^ profiling the adult human hippocampus, a subcortical brain region critical for episodic memory and spatial navigation.^53–55^ To ensure quality of the resulting cell types, we applied a stringent quality control criterion (number of expressed genes per nucleus > 600) and retained a total of 702,983 nuclei for downstream analyses (**Figure 6A**).

**Figure 6.**
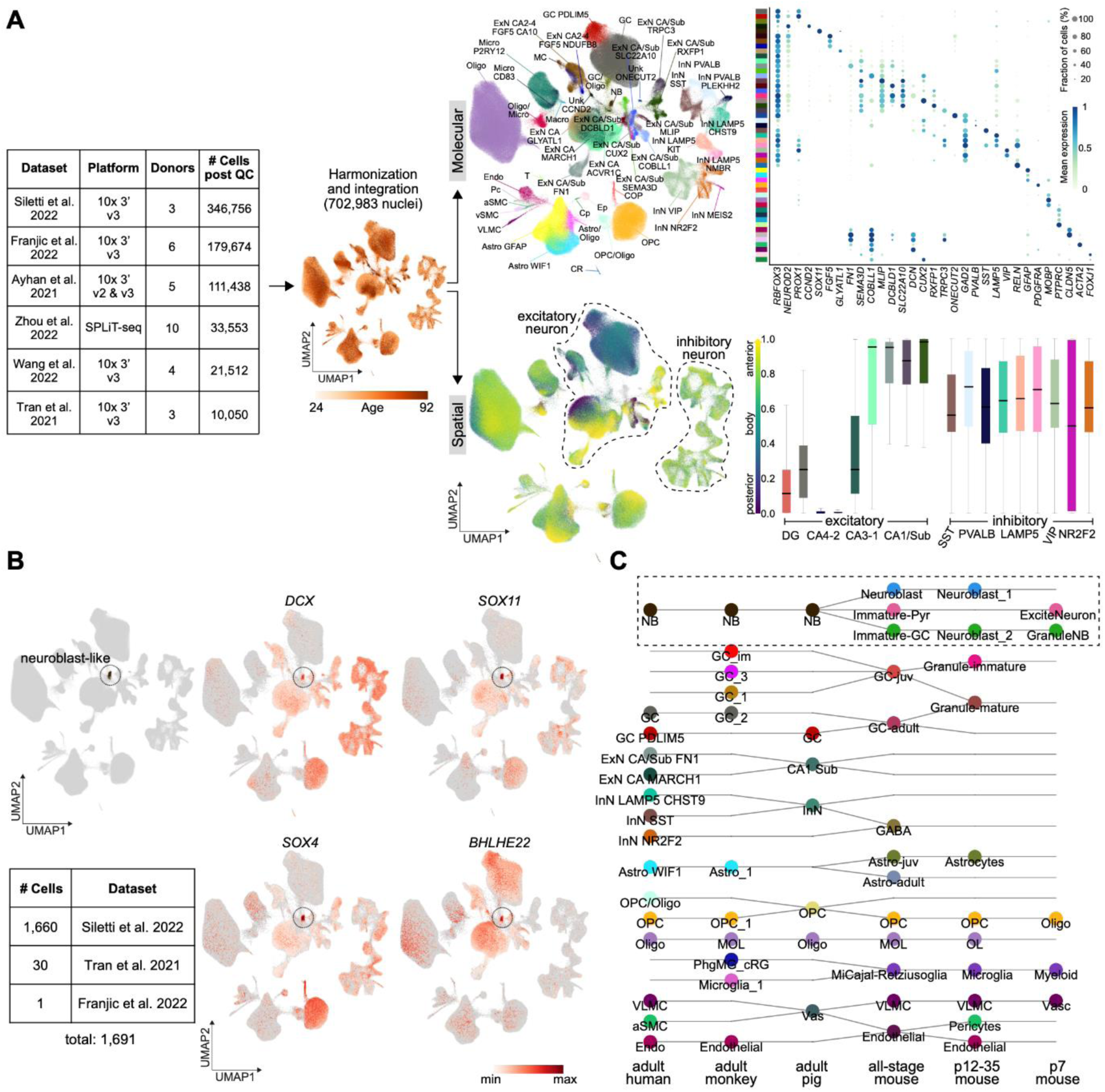
CellHint assembles a highly integrated adult human hippocampus atlas. (A) Left: the summary table showing the statistics of the six datasets used to assemble an adult human hippocampus atlas. UMAP to the right depicts the age distribution of these nuclei. Topright: the UMAP representation overlaid by the 51 cell types identified (left) and the dot plot showing expression of selected marker genes, with the color and size representing normalized gene expression and the percentage of cells expressing a given gene, respectively (right). Bottomright: the UMAP representation overlaid by the inferred anterior scores (left) and the box plots showing the distribution of these scores in selected excitatory and inhibitory neuron types, with the box frames extended from the lower quartile, median, to upper quartile (right). CA, cornu ammonis; Sub, subiculum; GC, granule cell; MC, mossy cell; ExN, excitatory neuron; InN, inhibitory neuron; CR, Cajal-Retzius cell; NB, neuroblast; Astro, astrocyte; OPC, oligodendrocyte progenitor cell; COP, committed OPC; Oligo, oligodendrocyte; Micro, microglia; Macro, macrophage; T, T cell; Endo, endothelial cell; Pc, pericyte; vSMC, venous smooth muscle cell; aSMC, arterial smooth muscle cell; VLMC, vascular and leptomeningeal cell; CP, choroid plexus; Ep, ependymal cell; Unk, unknown. (B) UMAP visualization highlighting the neuroblast-like cell population and its constituent datasets summarized in the table below. The four UMAP plots to the right are overlaid by expression of canonical markers of neuroblasts or immature neurons. (C) Harmonization graph showing cell type alignments for selected cell types from six datasets of three species. Neuroblasts and immature neurons are marked in a dashed rectangle. The datasets are, from left to right, integrated adult human data, adult macaque hippocampus,^60^ adult pig hippocampus,^48^ all-stage mouse hippocampus,^59^ mouse hippocampus from postnatal days 12-35,^59^ and mouse hippocampus from postnatal day 7.^58^ See also Figure S8.

Using CellHint, we integrated nuclei from 31 human donors which spanned ages from 24 to 92 years and exhibited a minimal batch effect (**Figure S8A**). Subsequently, we built on the CellHint output and added unsupervised clustering and in-depth manual curation (**Figure S8B**). This resulted in 51 cell types including 22 excitatory and 9 inhibitory neuron types, as well as diverse non-neuronal cell types constituting the hippocampal niche (**Figure 6A**). Cell types from the adult human hippocampus are relatively understudied as compared to those from the neocortex, with fewer phenotypic and genetic analyses in the literature. We thus utilized the transcriptomic features of the clusters to annotate them into molecularly defined populations (**Figure 6A**). This analysis revealed a marked cellular heterogeneity in the hippocampus, in particular for the excitatory neuron types from the cornu ammonis (CA) and subiculum. Here, we found a heterogeneous pool of excitatory neurons including both continuous and discrete cell states with characteristic molecular patterns (**Figure S8C**). To further enhance the interpretation of these cell types, we superimposed spatial information onto them, as the hippocampus has long been known to harbor a molecular and functional gradient along its longitudinal axis.^56, 57^ Specifically, we leveraged the datasets of Ayhan et al.^49^ and Siletti et al.^47^ where anterior-posterior and head-body-tail dissection information was available, respectively, and then inferred the anterior and posterior scores for cells in the remaining datasets (**Figure 6A; STAR Methods**). Through this, we reconstructed a global spatial map for the 51 cell types.

Among them, inhibitory neuron types did not display a significant bias towards anterior or posterior locations, indicating the lack of longitudinal variations. In contrast, the other cell types displayed varying levels of anteroposterior organizations along the hippocampal long axis. The most remarkable gradient was found in the excitatory neuron types, where the anterior scores gradually increased from the dentate gyrus, CA4-1, to the subiculum as expected, confirming the utility of our approach (**Figure 6A**). Therefore, on the basis of the molecular and inferred spatial information, we charted a detailed adult human hippocampus map to help interpret the cell types.

Interestingly, we detected a neuroblast-like population in this integrated atlas, indicating the possible presence of adult-born neurons (**Figure 6B**). Neurogenic trajectories have previously been detected at single-cell resolution in the adult hippocampus of mammalian species such as macaques, pigs, and mice;^48, 51, 58–60^ however, detection of human adult hippocampal neurogenesis from single-cell transcriptomics still remains controversial.^61^ Though human immature granule cells have been reported in two recent single-cell studies, they did not form a distinct cluster, being either scattered within the granule cell cluster and only identifiable by machine learning-based cell scoring^50^ or being at the border of the granule cells and thus of low confidence.^51^ In our atlas which integrated over 700,000 high-quality hippocampal cells, we located a clear neuroblast-like population. These cells were excitatory (*SLC17A7*+*GAD2*-), precluding the possible contamination of early inhibitory neurons from the adjacent neurogenic olfactory epithelium (**Figure S8D**). They were located between granule cells and non-dentate-gyrus excitatory neurons in the UMAP space. Multiple neuroblast-related biomarkers were overrepresented in these cells, including *DCX*, *SOX11*, *SOX4*, and *BHLHE22* (**Figure 6B**), as well as other markers characteristic of neuroblasts or immature neurons (e.g., *NNAT*, *STMN1*/*2*/*4*, *DPYSL3*, and *TUBB3*) (**Figure S8D**). Consistent with this, Gene Ontology analysis revealed the functional enrichment of this population in brain and axon development, regulation of cell development, protein maturation, and neuron projection development, as well as BMP signaling pathway and synapse structure or activity (**Figure S8E**). This population was composed of 1,691 cells from three datasets (**Figure 6B**). Among them, one single cell was from Franjic et al.,^48^ coinciding with only one neuroblast identified by the original study. 30 cells were contributed by the “Excit_H” cell type (30 out of 33) in Tran et al.^52^ Re-examination of “Excit_H” showed that it was highly intermingled with various neuronal populations and expressed a number of neuroblast markers (**Figure S8F**). The remaining 1,660 cells were mainly from the cluster 401 in Siletti et al., a cluster mostly dissected from uncal CA1-3.^47^

We next mapped cell types from this atlas onto cells from the hippocampus of adult *Macaca fascicularis* which was reported to have a complete repertoire of newborn neuronal types.^60^ The result showed that macaque neuroblasts mainly aligned with the human neuroblasts identified here, and to a lesser extent, with the mature human granule cells (**Figure S8G**). To further investigate their presence across species, we used CellHint to harmonize five additional hippocampus datasets from three mammalian species (macaque,^60^ pig,^48^ and mouse^58, 59^) (**Figures 6C and S8H)**. This analysis demonstrated the correspondence of the neuroblast population across human, macaque, and pig, with a further subdivision into three different immature excitatory neuron subtypes in the mouse hippocampus. This underscores the possible identity of these human cells as neuroblasts or immature neurons (**Figure 6C**).

### A multi-organ reference map for cell harmonization, integration, and classification

Integrating single-cell resources across the community, a current focus of the HCA consortium, is often laborious. Taking advantage of its highly automated workflow from harmonization to integration, we anticipate that CellHint could accelerate this process. We thus employed CellHint to build multi-organ reference maps following three steps: i) manual collection of public organ-focused datasets and their associated meta-data, ii) automatic cell type harmonization and integration by CellHint, iii) curation and finalization of the derived cell atlases (**Figure 7A**).

We compiled a total of 38 single-cell and single-nucleus transcriptomics datasets profiling 3,694,864 cells from 369 adult human donors in 12 tissues and organs, including the blood, bone marrow, heart, hippocampus, intestine, kidney, liver, lung, lymph node, pancreas, skeletal muscle, and spleen (**Table S2**). After applying the automated pipeline in CellHint, we generated 12 standardized organ reference atlases, available at https://www.celltypist.org/organs. All cells in these atlases were reannotated into the harmonized groups inferred by CellHint, which after further manual inspection and categorization were finally curated into biologically meaningful cell type labels. Based on this resource, we offered three complementary functionalities for the single-cell community (**Figure 7B**). Firstly, we provided a two-dimensional space (UMAP) onto which cells integrated across datasets were projected, enabling inspection of cell types and gene expression. Secondly, to demonstrate how cell types were harmonized and transcriptomic connections were established across datasets, we visualized the cell type alignments in the form of a harmonization graph or cell type hierarchy, referred to as the “tree plot” in CellHint. Thirdly, we leveraged the label transfer framework in CellTypist, a light-weight cell annotation algorithm based on logistic regression,^8^ to train a suite of machine learning classifiers. Thus, we provided 48 dataset-specific and 12 organ-specific models to achieve automatic cell type classifications in a broad tissue context.

## Discussion

Although the single-cell community has been using different batch correction methods to integrate cells, information about independently annotated cell types across datasets, a valuable part of single-cell genomics, has not been fully exploited. To benefit from annotations for cross-dataset integration, we developed workflows for cell type harmonization and integration, combined them into the CellHint infrastructure, and employed this pipeline to address challenges in developing cell type standards.

The PCT algorithm, along with other critical steps and optimizations, endows CellHint with scalability, speed, and accuracy (**Table S1**). We examined the performance of CellHint on five heterogeneous immune datasets, and found consistency between cell types harmonized by CellHint and those curated manually. In CellHint, we adopt the harmonization graph to iteratively redefine and reannotate cell types without losing relationships between any pairs of datasets. Within this graph, CellHint defines the relations between cell types using predefined symbols such as “=”, “∈” and “∋”, which are shown to cover the vast majority of cell type relationships found across studies and organs. This approach overcomes the challenge in multi-dataset alignment and dataset order instability, and generates consistent cell type hierarchies.

Reconstructing cell type hierarchies is a long-standing theme in cell biology. Whereas cell type relationships are imprinted in their molecular traits, transcriptome-based cell groupings (e.g., via unsupervised hierarchical clustering) do not necessarily mirror developmental history or biology. Expert knowledge is needed to group or segregate cell types regardless of their transcriptomic similarities. We speculate that this hard-earned legacy knowledge is often already incorporated into manual annotations, and our approach now allows this information to be integrated across multiple datasets to approximate the underlying biological hierarchies. CellHint represents an effort towards this direction and lays the foundation for future work aiming to incorporate knowledge into data-driven hierarchies (and vice versa). Compared to an earlier work which used hierarchical progressive learning to build cell type hierarchies,^62^ CellHint is able to mitigate the differences across datasets, thus discovering cell type relationships in a batch-insensitive manner. In practice, harmonization of cell types across datasets allows for selection of optimal or suboptimal clustering resolution that can best mirror the resulting cell subtype division schema underlying the cell hierarchy.

Application of CellHint to additional datasets further demonstrates its potential to unveil molecular and cellular variations in both physiology and pathology. Since cells from independent laboratories are harmonized, the resulting information can be exploited to put each cell into different contexts beyond its original study. This leads to consolidation of well-defined cell types and identification of novel cell states, as well as alignment of the underlying gene expression signatures and their altered characteristics in disease. In the four diseased lung datasets, CellHint was able to uncover pathological cell types and states that could only be identified when multiple datasets were harmonized and greater resolution was achieved. In the six datasets profiling the adult human hippocampus, CellHint discovered a potential neuroblast population. It should be noted that this population was probably localized in CA instead of dentate gyrus, thereby warranting further investigation.

For data integration, CellHint uses an annotation-aware algorithm to supervise the construction of neighborhood graphs. Influence of cell annotation on the data structure can range from forcibly merging the same cell types to a more lenient cell grouping. Adjusting the stringency of cell type annotations provides an effective way of integrating the cell type hierarchy into the data integration process. We show that CellHint is able to integrate diverse datasets to alleviate their batch effects and maintain the cell type relationships across datasets. Having a consistent database and search architecture to quantitatively compare cell types is critical to the single-cell field.^63–65^ The seamless integration of the two functionalities (harmonization and integration) in CellHint allows us to semi-automate the atlas building effort. We harmonized ∼250 high-quality cell types from 12 human tissues, providing a multi-organ resource that allows information of cell type harmonization, integration, and classification to be simultaneously inspected and queried. We also anticipate that this highly structured CellHint architecture, together with the multi-organ landmark resource, will aid in the assembly of diverse single-cell resources, facilitate the propagation of cell knowledge across communities, and improve the interpretability of organ-to-organ and health-to-disease cell relationships.

### Limitations of the study

Here we developed CellHint to harmonize and integrate cell types across different single-cell datasets. The unit of analysis is the collective identity referred to by a cluster of cells, namely, cell annotation. In this regard, spot-based assays such as spatial transcriptomics often do not fit in our pipeline, especially for spots that contain varying cell type mixtures. Therefore, CellHint is currently focused on cell-based genomics assays in order to provide the community with high-quality annotations at the cell and cell type levels. Moreover, CellHint employs a harmonization graph to visualize different cell annotations across datasets. As a result, the structure of the resulting hierarchy depends on the quality of cell types, in particular, the difference in annotation resolution across datasets. In addition, due to constraints on representation in two dimensions, cell types with complex one-to-one and one-to-many alignments may be overlaid in the resulting tree plot. Though these cross-connections are partially resolvable through visualization with a different order of datasets, three-dimensional tree plot or other forms of visualizations may be further avenues to pursue. Another limitation of CellHint is that in datasets with highly abundant ambient RNAs, not all cell types belonging to the same category can be well matched across datasets (**Figure S2F**). The development of methods to distinguish these artifacts from real transcriptomic deviations is a topic of interest for further developing CellHint.

## Acknowledgments

We thank D. Sumanaweera and K. Polanski for feedback on the CellHint algorithm; A. Cujba, E. Madissoon, and other members of the Teichmann lab for helpful discussions; A. Maartens and R. Vento-Tormo for editing and commenting on the manuscript. This research was funded in whole, or in part, by the Wellcome Trust Grant 206194 and 220540/Z/20/A. This work is also funded by the Technology Vision 2.0 under Turing project code R-TEC-001 which in turn is funded by the Engineering and Physical Sciences Research Council (EPSRC). This publication is part of the Human Cell Atlas (www.humancellatlas.org/publications/).

## Author contributions

C.X. and S.A.T. conceived the project. C.X. developed algorithms, collected data and performed all analyses. C.X., S.W., L.J., B.J.S., R.H., and P.H. contributed to the immune cell annotation in the CellTypist database. C.X. and M.P. structured the database and web portal. C.X. and S.A.T. wrote the manuscript with modifications from K.M..

## Declaration of interests

S.A.T. is a remunerated member of the Scientific Advisory Board, Foresite Labs, and Qiagen. She is co-founder and equity holder of Transition Bio.

## Inclusion and diversity

We support inclusive, diverse, and equitable conduct of research.

---

**Figure S1.**
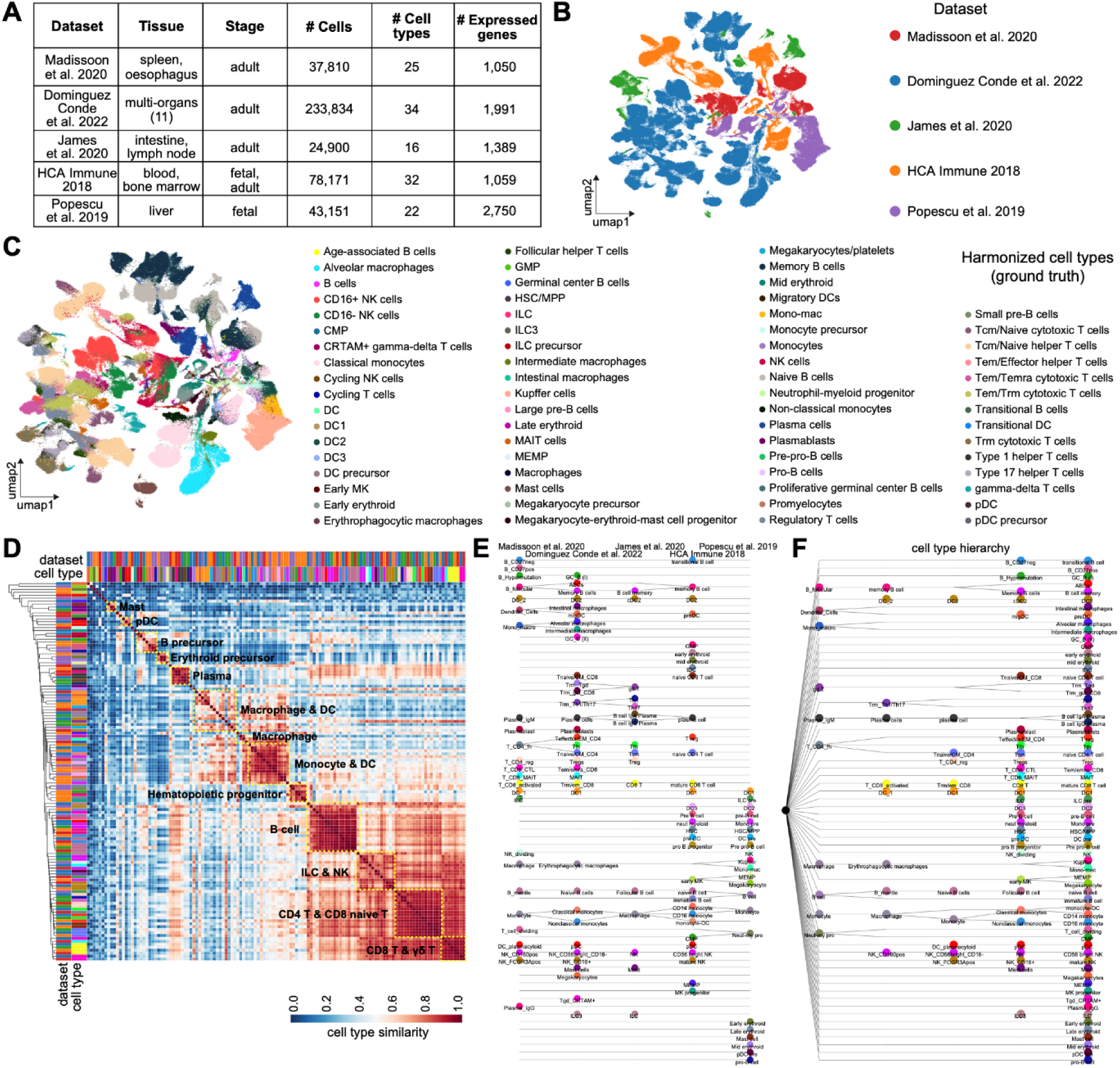
Harmonization of cell types in the five immune datasets, related to Figure 2. (A) Table showing the summary information (tissue, developmental stage, numbers of cells, cell types and expressed genes) of each dataset. (B) UMAP visualization of all cells colored by the dataset of origin, on the basis of a canonical PCA-based single-cell pipeline. (C) As in (B), but colored by the manually curated cell types. (D) Heat map displaying the unsupervised hierarchical clustering of cell types based on their inferred similarities by CellHint. Cell types are colored as in (C), and clustered into cell compartments marked in the plot. (E) Harmonization graph showing cell type relationships across five immune datasets shown on top. Cell type labels are colored according to their low-hierarchy alignments. (F) Cell type hierarchy reorganized from (E).

**Figure S2.**
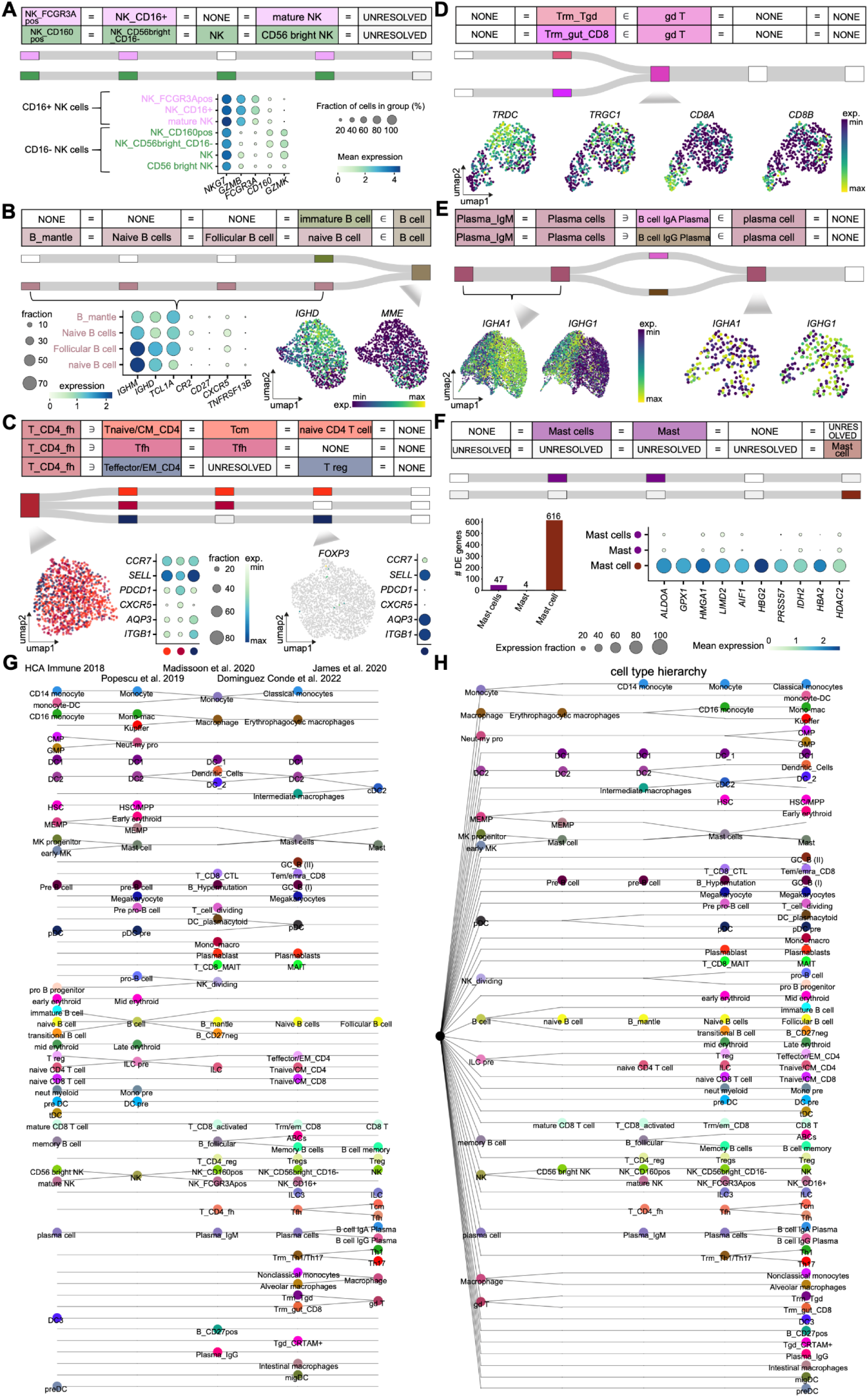
Examples of immune cell types and subtypes harmonized by CellHint, related to Figure 2. (A) Upper: part of the harmonization table containing seven natural killer (NK) cell populations, as well as their alignments visualized as a Sankey diagram. The order of datasets is, from left to right, Madissoon et al.,^23^ Dominguez Conde et al.,^8^ James et al.,^25^ HCA Immune 2018, to Popescu et al.,^24^ and applies to the rest of panels. Lower: dot plot showing expression of marker genes across the seven cell populations corresponding to two subtypes harmonized by CellHint. Color of the dot represents normalized gene expression, and size represents the percentage of cells expressing a given gene. (B) As in (A), but for the six B cell-related populations, including a “B cell” population that is harmonized into a spectrum of immature and naive B cells. (C) As in (A), but for the eight CD4+ T cell-related populations, including a “T_CD4_fh” population that is harmonized into three transcriptionally similar subtypes and a “T reg” population that was mis-annotated as demonstrated by the lack of *FOXP3* expression and is actually an effector memory CD4+ T cell subtype. (D) As in (A), but for the three cell populations transcriptomically similar with γδ T cells in the gut, including the “gd T” population that contains a hidden gut-resident CD8+ T cell subtype. (E) As in (A), but for the five plasma cell-related populations, including plasma cells from three datasets that are harmonized into IgA and IgG plasma subtypes. (F) As in (A), but for the three mast cell-related populations, including a “Mast cell” population highly abundant in ambient RNAs (and also with more differentially expressed genes) and failing to match other mast cells. (G) Harmonization graph showing cell type relationships across five immune datasets shown on top, with a different order of datasets as compared to Figure 2A and Figure S1E. Cell type labels are colored according to their low-hierarchy alignments. (H) Cell type hierarchy reorganized from (G).

**Figure S3.**
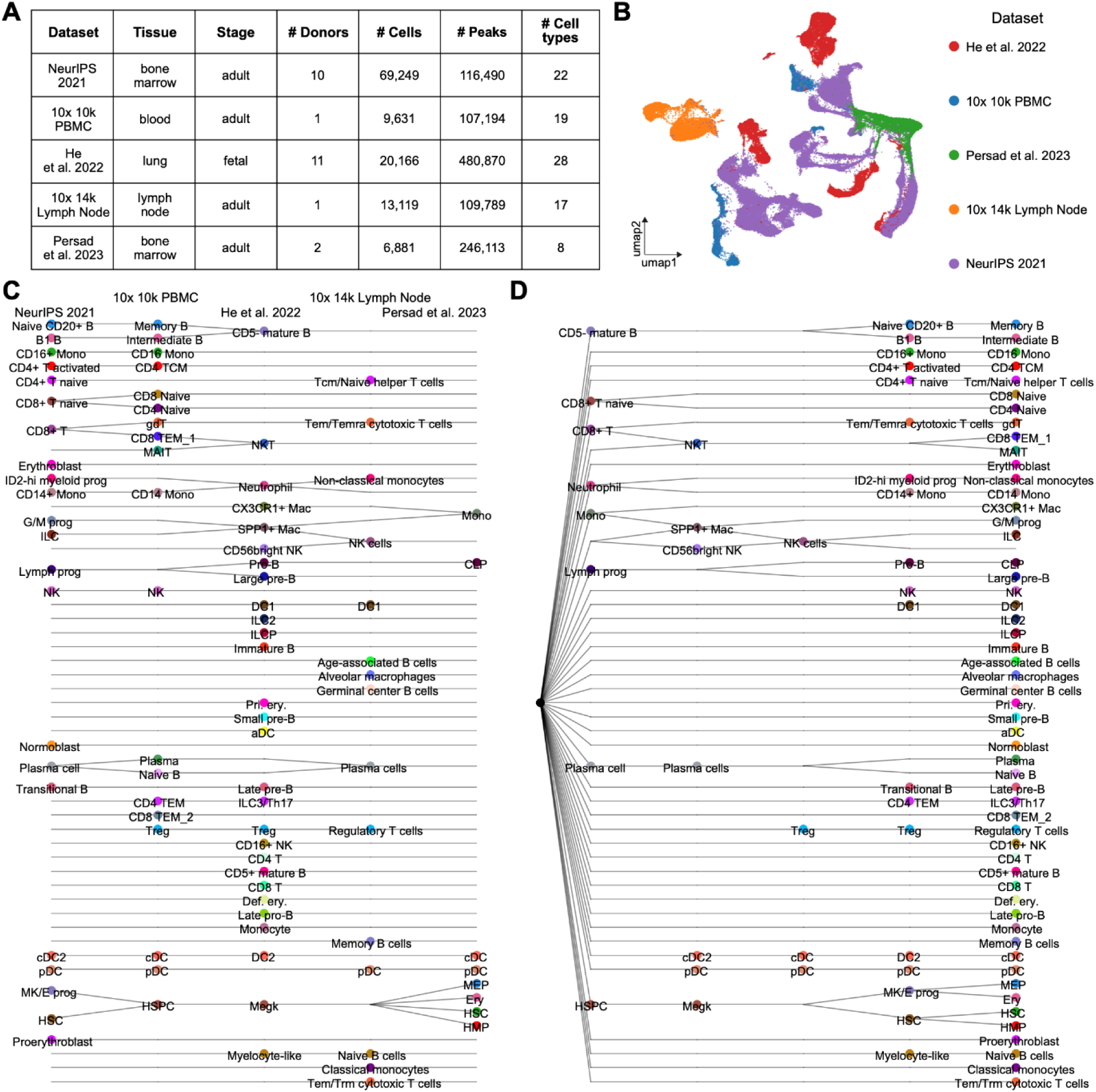
Harmonization of immune cell types from five chromatin accessibility datasets, related to Figure 2. (A) Table showing the information summarized for each dataset (tissue, developmental stage, numbers of donors, cells, peaks, and cell types). (B) UMAP visualization of all cells colored by the dataset of origin. (C) Harmonization graph showing cell type relationships across the five datasets shown on top. Cell type labels are colored according to their low-hierarchy alignments. (D) Cell type hierarchy reorganized from (C).

**Figure S4.**
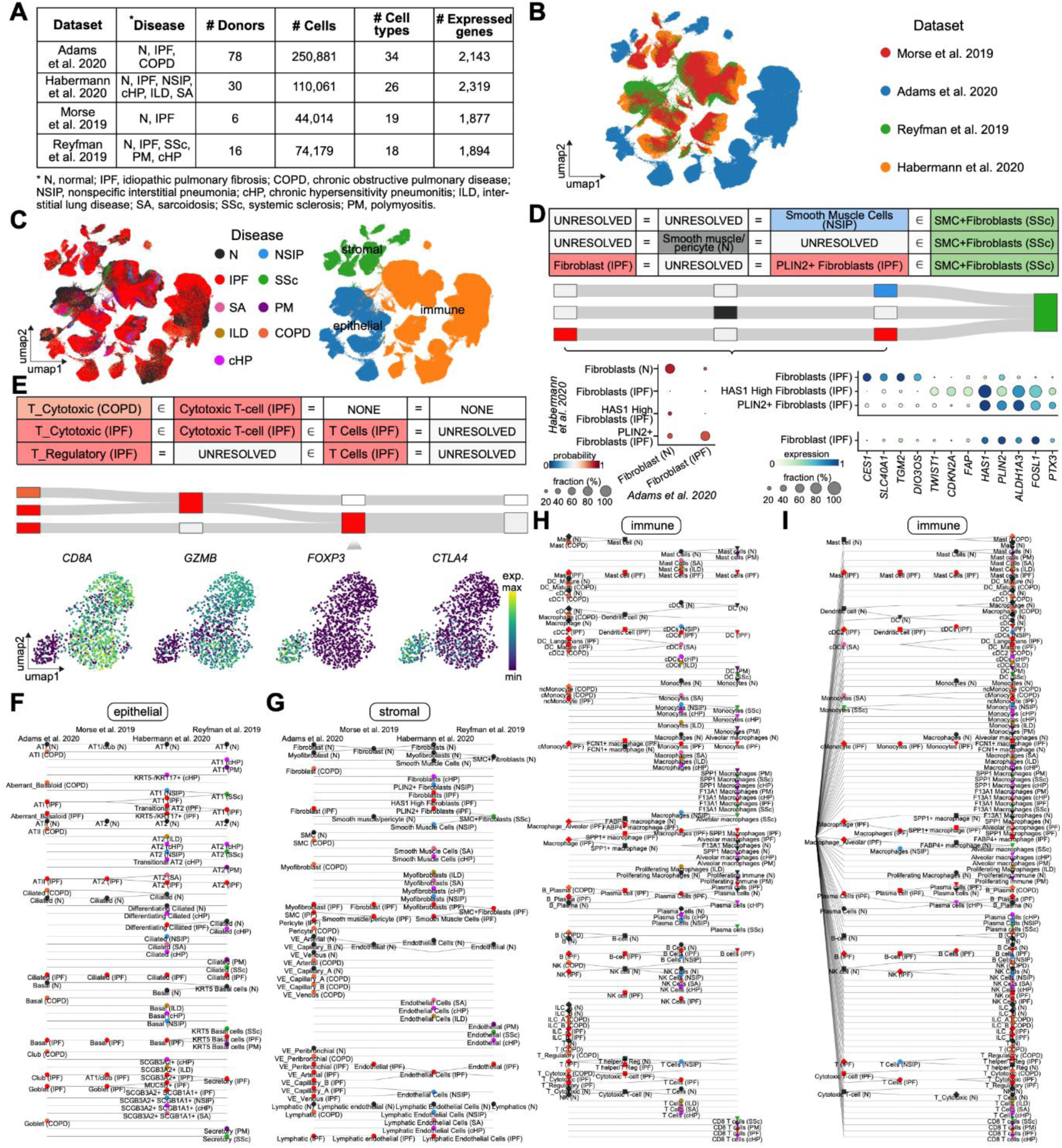
Harmonization of cell types in the four diseased lung datasets, related to Figure 3. (A) Table showing the summary information (disease, donor, numbers of cells, cell types and expressed genes) of each dataset. Abbreviations of diseases are summarized at the bottom of the table. (B) UMAP visualization of all cells colored by the dataset of origin, on the basis of a canonical PCA-based single-cell pipeline. (C) As in (B), but colored by the normal and eight disease conditions (left) and the three cell compartments (right). (D) Upper: part of the harmonization table containing fibroblasts and smooth muscle cells, as well as their alignments visualized as a Sankey diagram. The order of datasets is, from left to right, Adams et al.,^33^ Morse et al.,^34^ Habermann et al.,^35^ to Reyfman et al.,^36^ and applies to (E). Lower: scatter plot showing cell type label transfer from the third dataset to the first dataset, highlighting a correspondence between “PLIN2+ Fibroblasts” and “Fibroblast”. Color and size of the dot denote the mean prediction probability and cell type alignment fraction, respectively. Dot plot to the right shows expression of marker genes for the “Fibroblast” identified (bottom) as compared to those from the third dataset (top), with color and size of the dot representing normalized gene expression and percentage of cells expressing a given gene, respectively. (E) Upper: as in (D), but for a subset of T cell populations. Lower: UMAP representations overlaid by expression of marker genes of cytotoxic (*CD8A*+, *GZMB*+) and regulatory (*FOXP3*+, *CTLA4*+) T cell populations which together constitute the “T Cells” of the third dataset. (F) Harmonization graph showing epithelial cell type relationships across four lung datasets. Dataset names are shown on top. Color of the dot represents the normal or eight disease conditions for cell types. (G) As in (F), but for stromal cell types. (H) As in (F), but for immune cell types. (I) Cell type hierarchy reorganized from (H).

**Figure S5.**
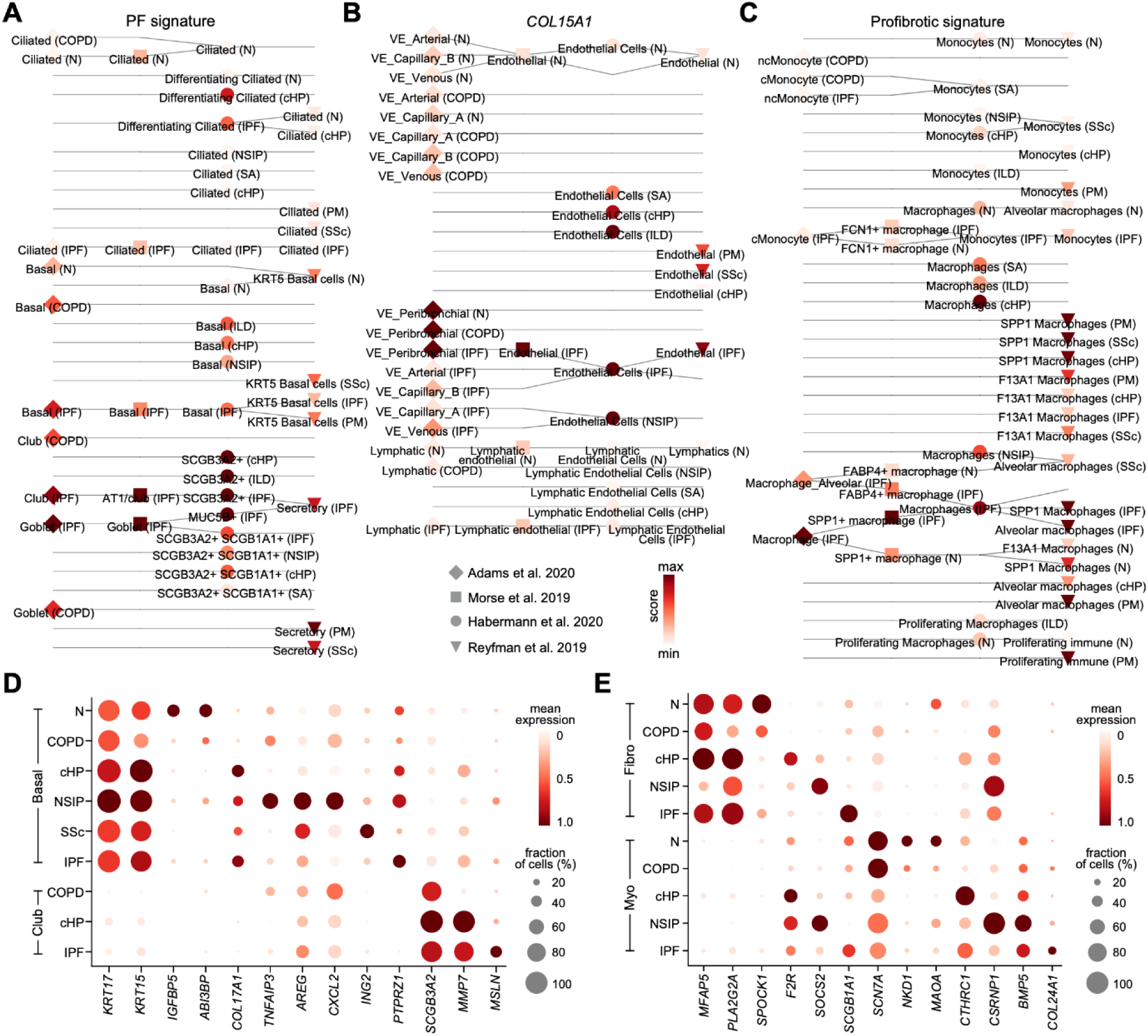
Gene signatures and molecular alterations across harmonized lung cell types, related to Figure 4. (A-C) Part of the harmonization graph that focuses on the pulmonary fibrosis (PF)-defining signature in airway epithelial cells (A), expression of *COL15A1* in endothelial cells (B), and the profibrotic signature in monocytes and macrophages (C). Color and shape of the dot represent the degree of a given signature and dataset of origin, respectively. (D) Dot plot showing expression of differentially expressed genes among basal and club cell types and among disease conditions, with the color and size representing normalized gene expression and the percentage of cells expressing a given gene, respectively. Cell types and diseases are grouped across datasets based on their alignments inferred by CellHint. (E) As in (D), but for the fibroblasts and myofibroblasts in the four datasets.

**Figure S6.**
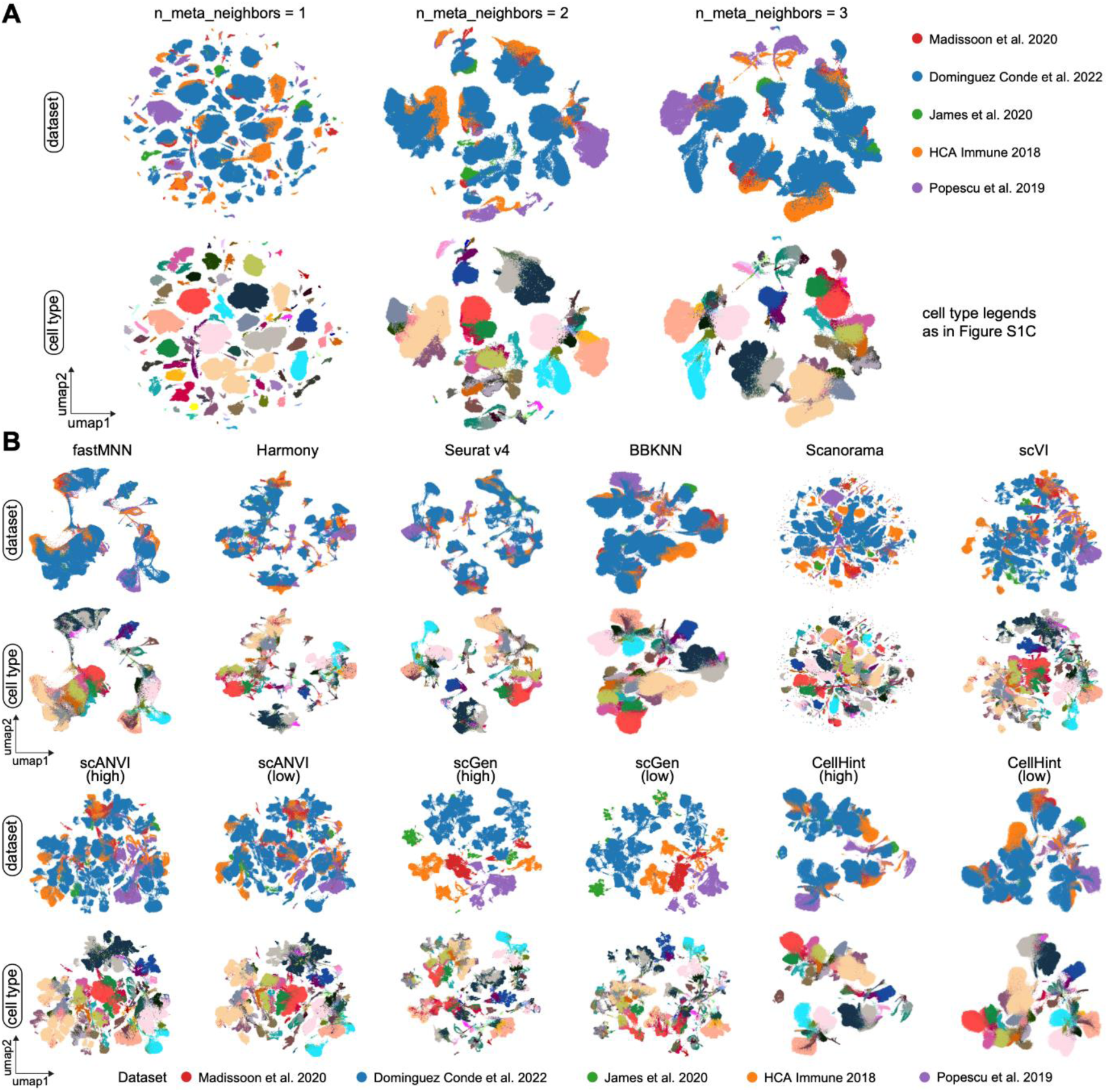
Integration of the five immune datasets using CellHint and other tools, related to Figure 5. (A) Impact of the number of cell type neighbors on transcriptomic structures. UMAP representations of cells from the five immune datasets are overlaid by the dataset of origin (top) and cell type (bottom). Transcriptomic structures are shown with different numbers of neighbors to anchor for each cell type when determining the group size of cell types: 1 (left), 2 (middle), and 3 (right). (B) Transcriptome organizations of immune cells integrated using different methods. UMAP representations of cells from the five immune datasets are integrated using six unsupervised (top two rows) and three supervised (bottom two rows) methods. For each method, cells are colored by the dataset of origin (upper) and the cell type (lower). For supervised methods, both high- and low-hierarchy annotations are used.

**Figure S7.**
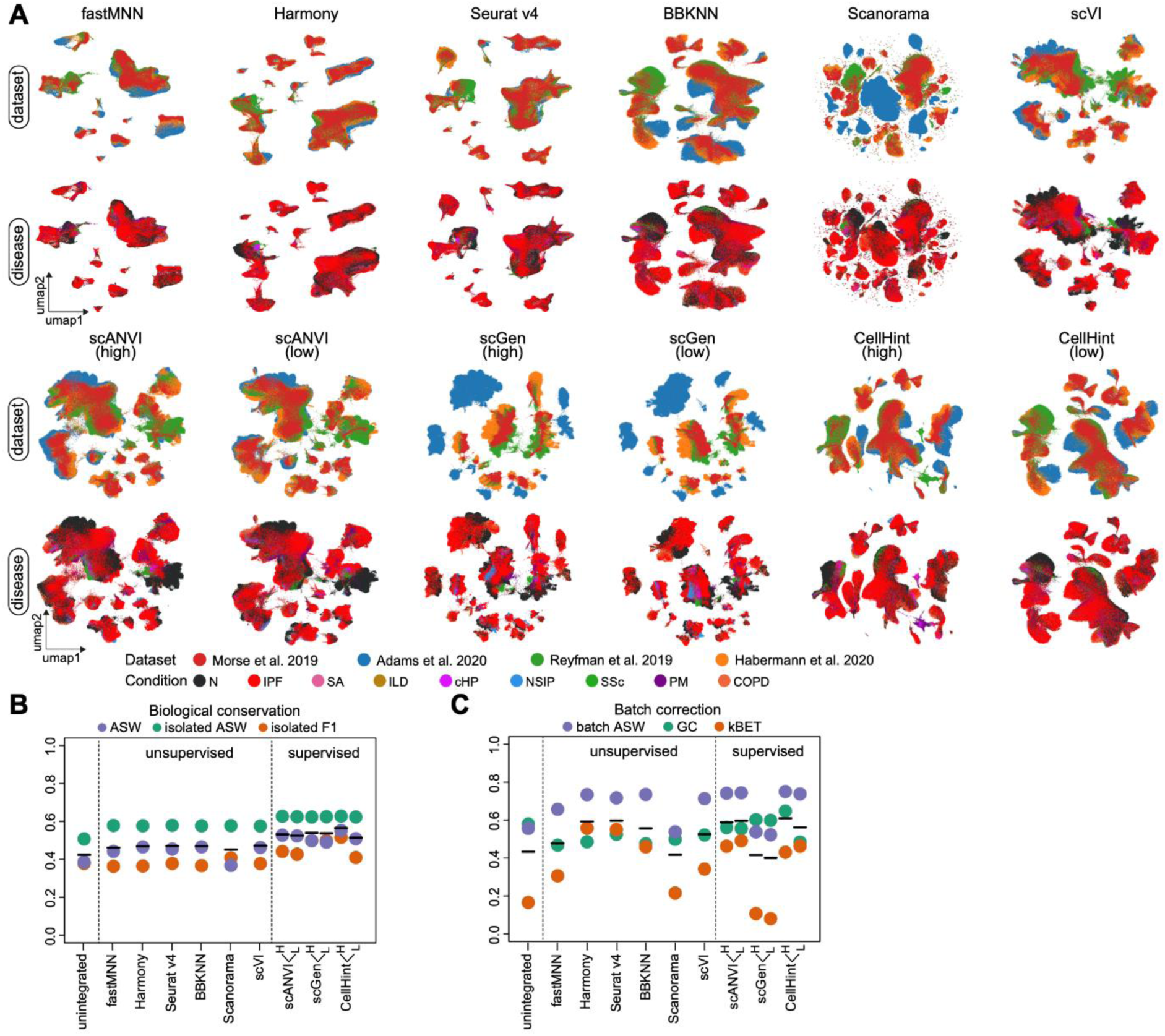
Integration of the four diseased lung datasets using CellHint and other tools, related to Figure 5. (A) Transcriptome organizations of lung cells integrated using different methods. UMAP representations of cells from the four lung datasets are integrated using six unsupervised (top two rows) and three supervised (bottom two rows) methods. For each method, cells are colored by the dataset of origin (upper) and the disease condition (lower). For supervised methods, both high- and low-hierarchy annotations are used. N, normal; IPF, idiopathic pulmonary fibrosis; COPD, chronic obstructive pulmonary disease; NSIP, nonspecific interstitial pneumonia; cHP, chronic hypersensitivity pneumonitis; ILD, interstitial lung disease; SA, sarcoidosis; SSc, systemic sclerosis; PM, polymyositis. (B) Assessment of biological conservation after lung data integration using the six unsupervised and three supervised methods. Names of tools are shown on the bottom. Both high- and low-hierarchy annotations are used for supervised methods. Horizontal black lines denote the averages of the integration metrics. ASW, average silhouette width; GC, graph connectivity; kBET, k-nearest-neighbor batch-effect test. (C) As in (B), bur for assessment of batch removal.

**Figure S8.**
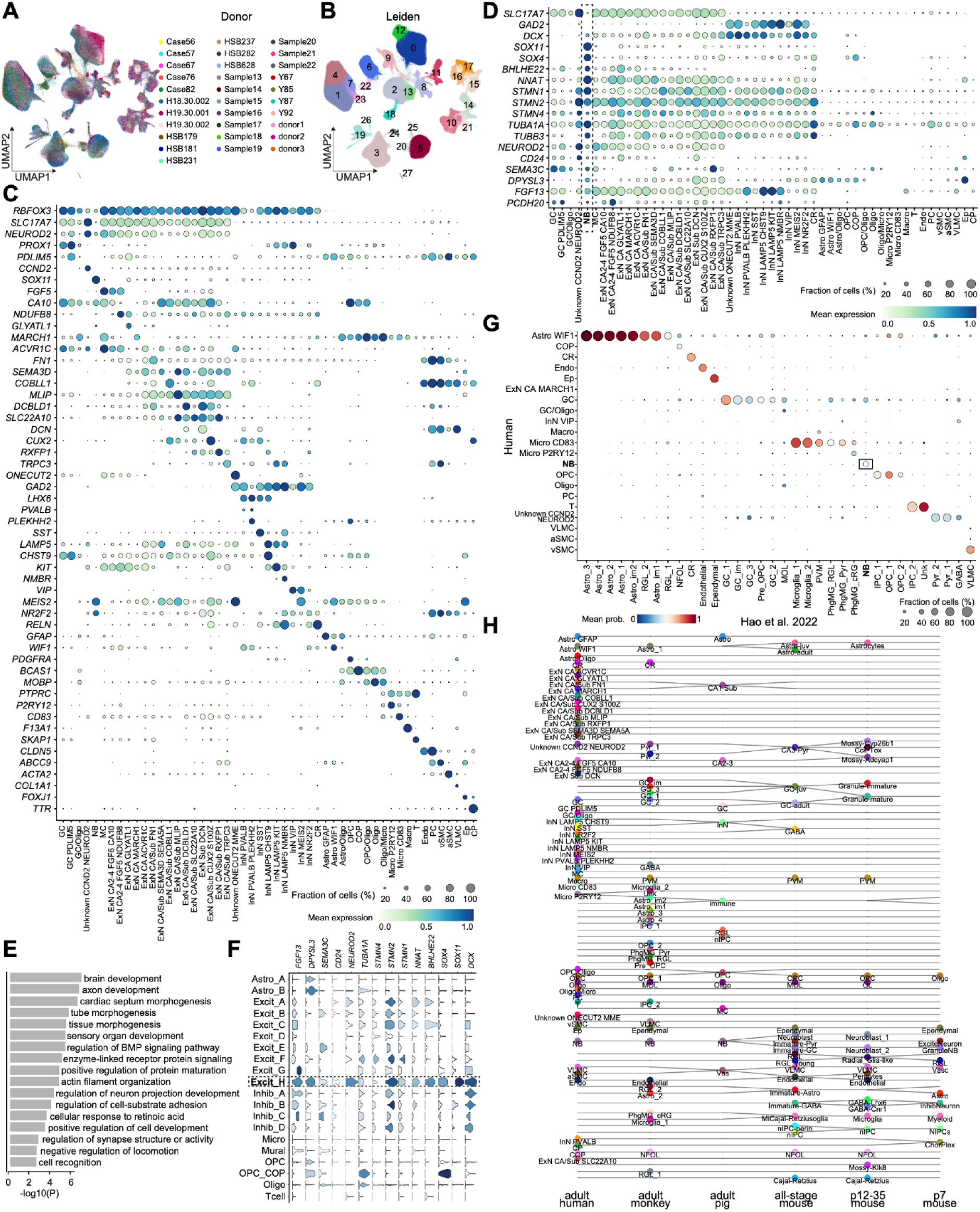
Cell types and their molecular features in the integrated adult human hippocampus atlas, related to Figure 6. (A) UMAP representation of all hippocampal nuclei colored by different donors. (B) As in (A), but colored by clusters derived from the Leiden clustering algorithm. (C) Dot plot showing expression of selected marker genes, with the color and size representing normalized gene expression and the percentage of cells expressing a given gene, respectively. (D) As in (C), but for markers related to the excitatory neuron (*SLC17A7*), inhibitory neuron (*GAD2*), and neuroblast- or immature neuron-related markers. (E) Bar plot depicting the *p*-values of the enriched Gene Ontology terms in the neuroblast-like population. (F) Stacked violin plots showing expression of neuroblast- or immature neuron-related markers in the “Excit_H” population in Tran et al.^52^ (G) Scatter plot showing cell type label transfer from the human dataset to the macaque dataset, highlighting a correspondence between “NB” in both datasets. Color and size of the dot denote the mean prediction probability and cell type alignment fraction, respectively. (H) Harmonization graph showing cell type alignments for all cell types from six hippocampus datasets of three species. The datasets are, from left to right, integrated adult human data, adult macaque hippocampus,^60^ adult pig hippocampus,^48^ all-stage mouse hippocampus,^59^ mouse hippocampus from postnatal days 12-35,^59^ and mouse hippocampus from postnatal day 7.^58^

## STAR Methods

## RESOURCE AVAILABILITY

### Lead contact

Further information and requests for resources should be directed to and will be fulfilled by the lead contact, Sarah A. Teichmann.

### Materials availability

This study did not generate new unique reagents.

### Data and code availability

● This paper analyzes existing, publicly available data. These accession numbers for the datasets are listed in the key resources table.
● All original code of CellHint has been deposited on GitHub and is publicly available. The link is listed in the key resources table.
● Any additional information required to reanalyze the data reported in this paper is available from the lead contact upon request.

## METHOD DETAILS

### The PCT algorithm

PCT is a decision tree-based algorithm for multi-target classification or regression analysis.^21, 66^ It is a top-down induced binary tree where each split results in two daughter nodes followed by iterative splits of the resulting daughter nodes until a stopping criterion is met. In a single-cell genomics context, PCT aims to recursively split all cells from a given dataset. Each split is executed by a specific expression cutoff from a given gene, and each resulting leaf corresponds to a derived cell cluster.

PCT is designed to account for the dependencies among target variables (here, attributes of cells), such that each cell within a given node or leaf is not described merely by one single attribute (e.g., cell identity or dissimilarity with one cell type) but by multiple attributes (e.g., membership or dissimilarities with multiple cell types). Each node or leaf has a prototype, defined as the eigen-pattern representative of all cells in this cluster, such as the average attribute profile of these cells. The distance between two prototypes from two daughter nodes respectively can be used to determine whether the split is necessary and what gene can be used for splitting.

For any query cell falling into one of the leaves in the reference tree according to its gene expression profile, its attribute profile will be predicted as the prototype of the leaf, thus taking into account the dependency among all attributes (i.e., all attributes of a reference cell are simultaneously used as a single vector when deriving the prototype). When the attributes of a cell are represented as a boolean vector denoting the likely membership of this cell with different cell types (e.g., defined at different levels in the hierarchy), PCT can be used for multi-level hierarchical classification with an interpretable probability output.^67, 68^ On the other hand, when the attributes of a cell are represented as a vector of dissimilarities with different cell types, PCT will be used as a multi-target regression algorithm. In CellHint, we adopted the latter as dissimilarity measures across cell types can be easily translated into cell type membership of a given cell. We thus define the transcriptional distance between two cell types as their dissimilarity (e.g., Euclidean distance at the PCA space or correlation distance at the gene expression space) if they come from the same batch or dataset, and as their inferred dissimilarity based on PCT if they come from two different batches or datasets. Another advantage is that due to the non-parametric nature of decision tree-related algorithms, data preprocessing steps that preserve the order of gene expression across cells have a marginal impact on induction of the trees; in other words, the trees built by PCT require little data preparation.

### CellHint workflow

A step-by-step explanation of the CellHint algorithm is detailed in the text below, which is also summarized in Table S1.

### Induction of the PCT-based tree

A direct calculation of the transcriptomic distance between two cells (or cell types) usually suffers from substantial batch effects when they originate from two different studies. To overcome this, for cell types across datasets or batches, CellHint leverages the PCT algorithm to predict the transcriptional dissimilarities instead of calculating them. The ultimate goal of this algorithm is to obtain a global matrix representing the distances between cells (rows) and cell types (columns) from all datasets. Since there is no ready-made PCT algorithm for use in the single-cell field, we adopt the commonly used tree structure, that is, parallel arrays from the Python scikit-learn package^69^ (version 0.24.1), and implement the PCT framework adapted for single-cell genomics.

A distance matrix 𝑑_𝑐,𝑐𝑡_ is first constructed for each dataset independently, where 𝑐 and 𝑐𝑡 represent the cells and cell types (defined as the cell centroids) in a given dataset, respectively. The general assumption here is that cells within one dataset are less impacted by batch effects in comparison to those across datasets. The metric of correlation distance (1 - Pearson’s 𝑟) or Euclidean distance is used to calculate this matrix based on the gene expression space or low-dimensional latent representations (e.g., PCA), respectively. For each cell 𝑐 (each row), this matrix records its dissimilarity profile across reference cell types.

We next define two key components of the clustering tree: distance measure and prototype.^21^ In CellHint, the dissimilarity between two cells (rows) in the matrix 𝑑 is assessed using the Euclidean distance, and the prototype 𝑝(𝑛) is defined as the dissimilarity profile averaged across cells that are involved during the tree induction of a given node 𝑛. In other words, cells with similar dissimilarity profiles tend to belong to a single node or leaf in the tree and their average profile is used as the prototype to represent the pattern of a query cell that falls into this node or leaf. The intra-node variance 𝜎^2^(𝑛) is defined as the sum of squared distances between each cell 𝑐 and the prototype 𝑝(𝑛):

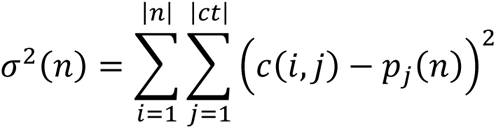

where 𝑖 enumerates each cell contained in the node 𝑛, 𝑗 enumerates each cell type in the reference dataset of interest, and 𝑐(𝑖, 𝑗) is the dissimilarity value for cell 𝑖 and cell type 𝑗. The benefit of such definition is that minimizing the intra-node variance during PCT is equivalent to minimizing the squared error when considering each 𝑐𝑡 independently and further summing them up across all cell types as defined in a classical multi-output regression tree (CART):

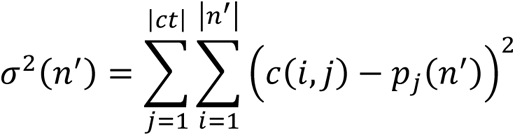

Also through this definition, during the top-down induction of the clustering tree, the node impurity (i.e., the degree of difference among cells in a given node) will be calculated in a similar way as CART. Specifically, since the impurity 𝐼(𝑛) for the node 𝑛 in CART is calculated as the mean squared error further averaged across targets (cell types), the node impurity for a PCT will be |𝑐𝑡| × 𝐼(𝑛) by treating the dissimilarity profile of each cell as a single response variable. The proxy impurity improvement is then quantified by summing the impurities of the two daughters of the node 𝑛 and adding a negative sign.

The clustering tree is split at each possible node by a “test” corresponding to an expression cutoff in a given gene that mostly separates cells within this node into two clusters (e.g., daughter nodes or leaves). By incorporating all above optimizations, a clustering tree is induced for each dataset, with a stringent requirement for node sizes (i.e., the number of cells to be greater than 20 which is adjustable in CellHint). Therefore, cells in a given dataset are recursively distributed along this tree with each branching event corresponding to a specific gene. However, the clustering tree is not exhaustively split to avoid overfitting. In the meantime, the resulting tree is not pruned by cost complexity pruning (CCP) as used by many tree-related algorithms, but by a more intuitive F-test as detailed below.

### Impurity reduction test and tree pruning

CellHint uses an F-test to determine whether the clustering tree needs to be further induced to a lower hierarchy for generating more leaves or cell clusters. Compared to CCP, this approach underscores the clustering tendency of similar cells which likely correspond to a homogeneous cell state but places less emphasis on the tree complexity (i.e., number of leaves or cell clusters generated by possible splits). The level of significance for a possible split at the node 𝑛 in the reference dataset can be assessed by an F-test:

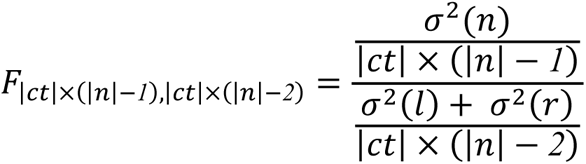

where 𝜎^2^(𝑛), 𝜎^2^(𝑙), and 𝜎^2^(𝑟) are the intra-cluster variances in the node, left daughter node, and right daughter node, respectively. Replacing all the three variances in this equation with the impurity values (𝐼(𝑛), 𝐼(𝑙), 𝐼(𝑟)) in CART allows us to infer the F-statistics in PCT as:

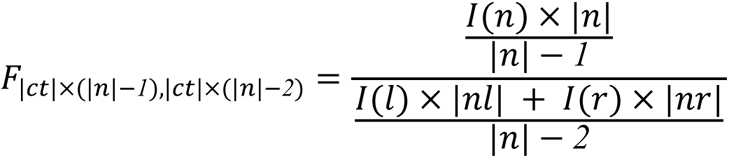

A *p*-value cutoff which defaults to 0.05 in CellHint is used to prune the node (and its recursive descendent groups) which has a small F-statistics.

### Prediction of dissimilarity profiles across datasets

The main aim of this section is to expand the cell-by-cell-type distance matrix in each reference seeding dataset to all cells (i.e., expanding rows) followed by expansion of cell types (i.e., expanding columns). Specifically, after a PCT is built for each dataset separately, it will be used to predict the cell-by-cell-type distance matrix 𝑑 for cells in the remaining datasets. Each query cell will be, according to its gene expression profile, assigned to one of the leaves in the clustering tree and predicted to be the prototype. The resulting distance matrices are then concatenated across cells from all datasets to form an expanded matrix 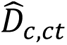 where 𝑐 covers cells from all datasets and 𝑐𝑡 represents the cell types from the reference dataset of interest. Prediction accuracy is calculated as the coefficient of determination (𝑅^2^) between the actual (𝐷_𝑐,𝑐𝑡_) and predicted 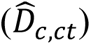dissimilarity profiles:

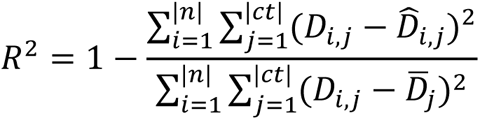

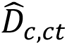 is next turned into a rank matrix starting from 1 to the number of elements in this matrix, and is further maximum-normalized to lie between 0 and 1. This is also the strategy adopted by MetaNeighbor.^70^ Finally, all 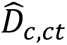 (each dataset has one 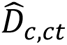) are concatenated across datasets to expand the repertoire of cell types, forming a global matrix 𝐺_𝑐,𝑐𝑡_ of dissimilarities among all cells (𝑐) and all cell types (𝑐𝑡). Of note, because a cell type is usually defined at the cluster level and theoretically no cluster is pure, CellHint can optionally filter out cells whose gene expression profiles do not correlate most with the cell type they belong to. This will speed up the run as only a subset of cells are used, but will render these filtered cells unharmonized in the downstream workflow (see the section below).

Meta-analysis at the cell type level is performed by averaging cells (i.e., rows) from the same cell type in the matrix 𝐺, leading to a cell-type-by-cell-type distance matrix among all cell types studied. This matrix is further transposed and averaged with the original matrix to form the final matrix that enables a consistent visualization given its symmetry.

### Cell type harmonization

Before undergoing the following stages, all cells are predicted into the most likely cell types in each dataset, on the basis of the global matrix 𝐺 mentioned above. Specifically, for each dataset, except for the cells which originate from this dataset, any query cell from the other datasets will be assigned one of the cell types that has the smallest distance with this query cell. We denote the result as the assignment matrix 𝐴 ∈ 𝑅^𝑐×𝑑^ in CellHint depicting how each cell 𝑐 (each row) is designated in each dataset 𝑑 (each column).

#### Stage I: initialization of the dataset core and definition of cell type relations

Two datasets are first selected, named as D1 and D2. Using their cross-prediction information in 𝐴, CellHint calculates two confusion matrices 𝑀*12* and 𝑀*21*, representing how cell types in D1 (rows) are assigned across cell types of D2 (columns), and vice versa. CellHint performs a pairwise alignment to form the dataset core for the two datasets with an iterative cell type removal procedure:

1. Initialize an empty harmonization table 𝐻𝑇 with three columns (D1, relation, D2)
2. Normalize each row of 𝑀*12* and 𝑀*21* to a sum of 1
3. Check whether novel cell types exist in D1 or D2. If yes, append them to 𝐻𝑇, remove them in both 𝑀*12* and 𝑀*21*, and go back to 2; if no, go to 4
4. Check whether one-to-one alignments exist. If yes, append them to 𝐻𝑇, remove them in 𝑀*12* and 𝑀*21*, row-normalize 𝑀*12* and 𝑀*21*, and go back to 4; if no, go to 5
5. Check whether one-to-many alignments exist. If yes, append them to 𝐻𝑇, remove them in 𝑀*12* and 𝑀*21*, row-normalize 𝑀*12* and 𝑀*21*, and go back to 5; if no, go to 6
6. Check whether many-to-one alignments exist. If yes, append them to 𝐻𝑇, remove them in 𝑀*12* and 𝑀*21*, row-normalize 𝑀*12* and 𝑀*21*, and go back to 6; if no, go to 7
7. Check whether any cell types are still maintained in 𝑀*12* and 𝑀*21* after 1-6. If yes, append them to 𝐻𝑇 as unharmonized, done; if no, done

From step 3 onwards, if 𝑀*12* or 𝑀*21* becomes an empty matrix in any step, the workflow will immediately end. A novel cell type C in D1 is appended as “C = NONE”; a novel cell type C in D2 is appended as “NONE = C”; an 1:1 alignment between C1 from D1 and C2 from D2 is appended as “C1 = C2”; an 1:N alignment between C1 from D1 and C2 and C3 from D2 is appended as “C1 ∋ C2” and “C1 ∋ C3”; a N:1 alignment between C2 and C3 from D1 and C1 from D2 is appended as “C2 ∈ C1” and “C3 ∈ C1”; an unharmonized cell type C from D1 is appended as “C = UNRESOLVED”; an unharmonized cell type C from D2 is appended as “UNRESOLVED = C”.

Definitions of the above seven categories are as follows: 1) Based on 𝑀*21*, if no cell types in D2 can align with a cell type C in D1 with more than *maximum_novel_percent* (default to 0.05 in CellHint), this is a novel cell type in D1. 2) Based on 𝑀*12*, if no cell types in D1 can align with a cell type C in D2 with more than *maximum_novel_percent* (default to 0.05 in CellHint), this is a novel cell type in D2. 3) In both 𝑀*12* and 𝑀*21*, if the alignments between C1 from D1 and C2 from D2 are greater than *minimum_unique_percent* (default to 0.5), plus that these alignments are not 1:N or N:1 (see 4 and 5), this will be an 1:1 match between C1 and C2. 4) If based on 𝑀*12*, one cell type C in D1 has more than two cell types aligned in D2 with a match proportion greater than *minimum_divide_percent* (default to 0.1), and these matched cell types in D2 have a back-match proportion greater than *minimum_unique_percent* (the same parameter used in 3), this will be an 1:N match. 5) If based on 𝑀*21*, one cell type C in D2 has more than two cell types aligned in D1 with a match proportion greater than *minimum_divide_percent*, and these matched cell types in D1 have a back-match proportion greater than *minimum_unique_percent*, this will be a N:1 match. 6) Based on 𝑀*12*, if after all these categorizations, a cell type in D1 remains unharmonized (at this time, 𝑀*12* should have no columns available), this will be an unharmonized cell type in D1. 7) Based on 𝑀*21*, if after all these categorizations, a cell type in D2 remains unharmonized, this will be an unharmonized cell type in D2. Of note, during multi-dataset (>=3) alignments, all three cutoffs used above are dynamically chosen to find the best alignments during iterations (see stage IV).

#### Stage II: combination of datasets and cell type renaming

The aim of this stage is to combine D1 and D2 into a new D1, allowing addition of a new D2 in stage III. As a first step, cells from D1 and D2 are combined, and the combined set of cells are called a new dataset D1. Next, CellHint reannotates the identities of these cells into cell types in the harmonization table 𝐻𝑇 obtained from stage I. Specifically, based on what cell type each cell is predicted to be in each dataset using the information from the assignment matrix 𝐴, each cell can be assigned one of the rows of 𝐻𝑇, or “UNASSIGNED” if this cell does not have a consistent alignment pattern existing in 𝐻𝑇. These “UNASSIGNED” cells are not considered in the following stages. Through this, CellHint obtains a “new” dataset with each resulting cell type re-annotated into a longer name corresponding to a given row of the harmonization table 𝐻𝑇.

#### Stage III: expansion of the dataset core and harmonization table

After the new D1 and associated annotations of cells in D1 are obtained, an additional dataset D2 (which is still named D2 because the earlier D1 and D2 have been combined and renamed to D1) is incorporated. The dataset core is expanded by aligning the D2 with D1 using a procedure below:

1. Form the 𝑀*12* matrix
2. Form the 𝑀*21* matrix
3. Perform the procedure in stage I with 𝑀*12* and 𝑀*21*, and obtain a new 𝐻𝑇
4. Disentangle 𝐻𝑇 to have multiple columns (one column per original dataset)
5. Perform the procedure in stage II with the new 𝐻𝑇, and form a new D1
6. Incorporate an additional dataset D2, and repeat stage III with the new D1

For step 1, 𝑀*12*, which represents the proportions of cell types in D2 (columns) assigned to cell types of D1 (rows), is formed by examination of the predicted D2 cell types for D1 cells, on the basis of the information from the assignment matrix 𝐴. For step 2, 𝑀*21* illustrates what cell types D2 cells are predicted to be in D1. Notably, since cells in D2 are predicted into cell types in D1 which have longer names and correspond to rows in 𝐻𝑇, not all cells in D2 can be annotated and are thus assigned “UNASSIGNED”, which are not used when generating 𝑀*21*.

After step 3, a new 𝐻𝑇 with three columns (D1, relation, and D2) is formed. Of note, the third column (D2) contains cell types from D2, whereas the first column (D1) contains cell types (rows) as defined in the earlier 𝐻𝑇. This new 𝐻𝑇 will be expanded to have multiple columns with one dataset per column joined by a “relation” column, serving as input for step 5. When all datasets are iteratively incorporated, the final output will be a 𝐻𝑇 covering all cell types across all datasets.

#### Stage IV: tuning the optimal dataset order and cell reannotation

Although all pairwise predictions between datasets are considered by the above stages (information from the assignment matrix 𝐴 which has all cell types from all datasets), an optimal order of datasets is still useful with respect to grouping of similar datasets and visualization. CellHint uses a greedy strategy to determine the optimal order of datasets incorporated into the 𝐻𝑇. CellHint first calculates the similarity between any pair of datasets. Next, two datasets with the highest similarity are added in the first iteration of the harmonization pipeline, followed by addition of another dataset which has the highest total similarity with the datasets already added in the previous round. Here, the inter-dataset similarity is quantified by the numbers of alignments between two datasets (with adjustable weights of 2, 1, -1, and -2 for one-to-one, one-to-many, novel, and unharmonized matches, respectively) divided by the total number of cell types, denoting the degree of cell type share between the two given datasets.

During stage I to III, combinations of five *minimum_unique_percent* (0.4, 0.5, 0.6, 0.7, 0.8) and three *minimum_divide_percent* (0.1, 0.15, 0.2) are exhaustively tried to find the best combo, by choosing the parameters that result in the least alignments when a new dataset is added. This procedure is repeatedly conducted in each iteration. After the final 𝐻𝑇 is obtained, each cell will be reannotated to one of the cell types (rows) in 𝐻𝑇 based on the assignment matrix 𝐴.

#### Stage V: reorganization and visualization of 𝐻𝑇 and its cell type hierarchy

An interconnected (i.e., connections spanning multiple rows of 𝐻𝑇) group of cell types is further defined at the high hierarchy for 𝐻𝑇, with each group being independent of other groups. This group corresponds to a high-hierarchy cell annotation located by CellHint and each member (row) corresponds to a low-hierarchy cell annotation. For each cell type group, if it contains novel (“NONE”) or unharmonized (“UNRESOLVED”) representations, these blank labels will be renamed in order to distinguish the many blank labels in the 𝐻𝑇. Specifically, a blank element in the first column (dataset) will be searched against cell types from the other columns across the given row until a non-blank one is found and renamed to it. A blank element in other columns (except the first column) will be searched in the opposite direction and renamed to the first non-blank element in the given row. The newly formed 𝐻𝑇 forms the basis for the Sankey plot and the flat tree plot in CellHint.

Next, columns (datasets) of each cell type group will be reordered according to how many unique values each column has, including the renamed blank cell types. To disentangle the cross connections in 𝐻𝑇 for visualizations, rows of each group are further reordered to enable consistent alignments to lie at bottom of each group. This reordered 𝐻𝑇 is used to plot the cell type hierarchy by cross-linking unique cell types including the renamed blank cell types across datasets.

### Annotation-aware data integration

CellHint employs the approach of restricted neighborhood search to integrate cells from different datasets (or batches), a concept similar to that used by BBKNN.^16^ After cell type harmonization (or other procedures that can result in consistent cell type labels across datasets), all cells are placed into a common naming schema. In CellHint, for each cell belonging to a specific cell type, its nearest neighbors can only be found out of cells that belong to the same cell type. Since this strategy cannot produce a fully connected neighborhood graph and each cell type is strongly separated with other cell types, CellHint uses the cell type hierarchy and meta-neighbors to further expand the search space and preserve cell type spectrum. Specifically, the cell type hierarchy, especially the cell types grouped at the high hierarchical level, contains information about what cell types are transcriptionally similar. During the neighborhood search, each cell will search, in addition to the cell type it originates from, other cell types at the high hierarchy. Moreover, CellHint leverages the distance matrix among cell types (see the section “*Prediction of dissimilarity profiles across datasets*”) to find the nearest neighbors of cell types. For each cell type in each dataset, two nearest cell types will be located (the default parameter of n_meta_neighbors is three, meaning that two other cell types will be incorporated). These cell types will be next combined across datasets to form a final cell type group, against which neighbors of each cell will be searched.

Due to the presence of dataset-specific cell types, some cell type groups may not span all datasets. For these groups, the neighbors of a query cell will be contributed only by datasets (or batches) that qualify, through providing more cell neighbors per dataset. For example, if only three out of four datasets contain this cell type, then each dataset needs to provide five neighboring cells in order to achieve a total number of 15 neighbors for a given query cell. The resulting neighborhood graph is further trimmed to keep the top largest connectivities for each cell. To tackle the problem of partial annotation such as “UNASSIGNED” cells derived from the harmonization pipeline, CellHint incorporates these cells into every cell type group and thus expands the size of each group. Accordingly, for these unannotated cells, their search space will be among all cell type groups available.

### Analysis of five immune transcriptomics datasets

The five immune datasets were downloaded from the CellTypist database^8^ and combined into one cell-by-gene count matrix. After normalization and log-transformation (with a pseudocount of 1), the highly variable genes (HVGs) were selected using the function scanpy.pp.highly_variable_genes (batch_key is set to the dataset of origin) from Scanpy^71^ (version 1.7.2), followed by data scaling (scanpy.pp.scale) and principal component analysis (PCA) (scanpy.tl.pca). Cell type harmonization and cell reannotation were conducted by following the aforementioned pipeline in the Methods section “*Cell type harmonization*”, collectively wrapped as the function cellhint.harmonize in CellHint. The result was visualized as a tree plot or a cell type hierarchy using the function cellhint.treeplot, details of which can be found in the stage V of the Methods section “*Cell type harmonization*”. Meta-analyses of cell types across datasets were performed by averaging the cell-by-cell-type matrix 𝐺 across rows (cells) that belong to the same cell types. To confirm the existence of cell subtypes (e.g., ABCs) harmonized and detected by CellHint, associated cells (e.g., memory B cells) were extracted and re-clustered using a canonical PCA-based clustering approach. Marker genes used for inspection and validation of cell subtypes were from the CellTypist immune cell encyclopedia (https://www.celltypist.org/encyclopedia/Immune/v2).

### Analysis of five immune chromatin accessibility datasets

The five scATAC-seq and snATAC-seq datasets were downloaded from respective studies,^26–28^ including two public multiome (RNA + ATAC) datasets provided by 10x Genomics (download links can be found in the key resources table). We collected both the peak-by-cell count matrices and fragment files for these datasets, alongside their detailed cell type annotations. 119,046 immune and progenitor cells were extracted from these datasets for downstream analyses.

Due to the greatest number of cells in the NeurIPS 2021 dataset, we set the 116,490 peaks in this dataset as a common set of peaks for all datasets. Next, we quantified the Tn5 cut sites in each peak associated with a given cell using the FeatureMatrix function in Signac^72^ (version 1.10.0). After concatenating their count matrices, we performed feature selection (FindTopFeatures with min.cutoff set as 20), TF-IDF normalization (RunTFIDF), singular value decomposition (SVD) (RunSVD). The resulting normalized expression and low-dimensional SVD embeddings (excluding the first component highly correlated with the sequencing depth) were used as input for CellHint harmonization using the function cellhint.harmonize.

### Analysis of four diseased lung transcriptomics datasets

The four lung datasets were downloaded from respective studies.^33–36^ For each dataset, cells with fewer than 200 expressed genes were discarded. We next combined cell type annotations with disease conditions to define the statuses of cells in the four datasets, and further removed the cell status with a size fewer than 20 cells. Normalization, log-transformation, HVGs selection, scaling, and PCA were performed as in the Methods section “*Analysis of five immune transcriptomics datasets*”. The harmonization graph and cell type hierarchy were obtained using the function cellhint.harmonize in CellHint, and visualized with cellhint.treeplot. Label transfer analysis (e.g., between “Aberrant_Basaloid” and “KRT5-/KRT17+”) was conducted using the logistic regression framework in CellTypist based on the model training (celltypist.train) followed by cell type prediction (celltypist.annotate) with the default parameters. Existence of cell subtypes harmonized and revealed by CellHint was confirmed in a similar manner as that for the five immune transcriptomics datasets.

The following gene signatures were used. Pulmonary fibrosis: *CDKN2A*, *COL1A1*, *FN1*, *MMP7*, *MUC5B*, *SMAD3*, *ITGB6*, *GDF15*, and *EPHB2*; ECM: from the Gene Ontology (GO) term of “extracellular matrix structural constituent”; profibrotic signature: *SPP1*, *LIPA*, *SPARC*, *GPC4*, *PALLD*, *CTSK*, *MMP9*, and *CSF1*. Scores of gene signatures were calculated using the Scanpy function scanpy.tl.score_genes to account for background gene expression levels. For the analysis of molecular alterations, differential expression analysis was performed using the Scanpy function scanpy.tl.rank_genes_groups (with the parameter pts turned on to enable quantification of percentage of gene expression in a given cell type), based on the grouped cell types and diseases from the harmonization graph. Genes were selected with three criteria: a *p*-value less than 0.05, expression percent in the cell type of interest greater than 0.4, and expression percent in other cell types less than 0.2. A subset of genes with pathological relevance were selected and displayed ultimately.

### Analysis of six transcriptomics datasets of adult human hippocampus

The six datasets profiling the adult human hippocampus were downloaded from respective studies.^47–52^ We removed nuclei with the numbers of expressed genes fewer than 600 to only preserve high-quality nuclei. Top 3,500 HVGs were extracted from each study using the function scanpy.pp.highly_variable_genes (batch_key is set to the donor) from Scanpy, and the top 2,500 genes with the highest occurrence frequencies across datasets were finally selected. Normalization, log-transformation, data scaling, and PCA were performed as in the Methods section “*Analysis of five immune transcriptomics datasets*”. Due to a failure to collect annotation information from Wang et al.,^51^ cells from this dataset were labeled as “UNASSIGNED” which could be recognized by CellHint. Cell type harmonization and integration were conducted using CellHint, with the algorithmic details in the above sections. To further refine the structure of the neighborhood graph, after the cell types were manually curated (see below), we performed a second round of supervised integration with the new version of cell annotations. The Scanpy function scanpy.tl.umap was next utilized to generate UMAP embeddings.

Cell types were manually annotated by considering both the harmonization results and the unsupervised Leiden clustering. Both known markers of cell types and markers identified from differential expression analysis (using the function scanpy.tl.rank_genes_groups) were used to molecularly define the resulting cell types. To superimpose longitudinal information onto these cell types, we focused on two datasets which had anteroposterior^49^ and head-body-tail^47^ information, respectively. Specifically, for each cell type we trained two logistic regression models using CellTypist^8^ based on cells from the two datasets, respectively: one for anteroposterior classification and the other for head-body-tail classification. After applying these two models to all cells in the given cell type, for each cell we obtained the anterior and posterior scores from the first model and the head, body, and tail scores from the second model. The final anterior score of each cell was then defined as the weighted sum of anterior (1) and posterior (0) positions (for the first model), or the weighted sum of anterior (1), middle (0.5), and posterior (0) positions (for the second model), with the weights being the scores derived from the two models. The two scores of each cell were then averaged across the two models.

To map hippocampal cell types across species, we downloaded datasets from four studies,^48, 58–60^ which profiled the hippocampus of adult *Macaca fascicularis*, adult *Sus scrofa*, and *Mus musculus* in various stages. Gene orthologs between the human and each other species were downloaded from Ensembl BioMart (Ensembl Genes 110), and only one-to-one orthologs were used for all the downstream analyses. A total of 7,885 genes were ultimately kept. To transfer cell type labels from the human to macaque, the function celltypist.train in CellTypist was used to train a logistic regression-based classifier using human data, with the manual annotation being the label key and feature selection turned on. Cells from the macaque dataset were then predicted into human cell types using the function celltypist.annotate and visualized using celltypist.dotplot to illustrate the cell type correspondence between human and macaque. To align cell types across multiple species, we used the harmonization algorithm as detailed in the above sections to match cell types from all six datasets (human, macaque, pig, and three mouse datasets). Top 50 markers from the neuroblast-like population were determined using the Scanpy function scanpy.tl.rank_genes_groups (with the parameter pts turned on to enable quantification of percentage of gene expression in a given cell type). Genes were selected with three criteria: a *p*-value less than 0.05, expression percent in the neuroblast population greater than 0.4, and expression percent in other cell types less than 0.2. Gene set enrichment analysis was performed using Metascape.^73^

### Integration of datasets with different tools

The five immune datasets or four diseased lung datasets were integrated in an annotation-aware manner using CellHint, with the algorithmic details in the Methods section “*Annotation-aware data integration*”. The dataset of origin was considered as a batch confounder and the cell reannotation information from the harmonization pipeline was treated as a biological label key. The default parameters (e.g., n_meta_neighbors = 3) were used. The Scanpy function scanpy.tl.umap was further utilized to generate UMAP embeddings. We also used the following eight methods to integrate the datasets.

1) FastMNN: normalized expression data (HVGs only) was used. The data was split by the dataset of origin, and served as input for the wrapper function RunFastMNN in the R package Seurat^9^ (version 4.3.0). After this, neighbors were located using the function FindNeighbors (tag-value pairs: dims = 1:50, reduction = "mnn") and UMAP was generated with the function RunUMAP (tag-value pairs: dims = 1:50, reduction = "mnn").
2) Harmony: normalized and scaled expression data (HVGs only) with pre-calculated PCA coordinates were used. The wrapper function RunHarmony in Seurat was used with the dataset of origin being a batch confounder. After this, neighbors were located using the function FindNeighbors (tag-value pairs: dims = 1:50, reduction = "harmony") and UMAP was generated with the function RunUMAP (tag-value pairs: dims = 1:50, reduction = "harmony").
3) Seurat V4: normalized expression data (HVGs only) was used. The data was split by the dataset of origin, and for each dataset gene expression was scaled separately. Next, the function FindIntegrationAnchors (tag-value pairs: normalization.method = ’LogNormalize’, scale = FALSE, dims = 1:50) followed by IntegrateData (tag-value pairs: normalization.method = ’LogNormalize’, dims = 1:50) were used to generate a batch-corrected expression matrix. This matrix underwent a canonical pipeline of data scaling (ScaleData), PCA (RunPCA), neighborhood search (FindNeighbors), and UMAP generation (RunUMAP).
4) BBKNN: pre-calculated PCA coordinates were used. The function bbknn.bbknn was applied to the data with the dataset of origin being a batch confounder. The resulting neighborhood graph was used to generate UMAP embeddings with the function scanpy.tl.umap in Scanpy.
5) Scanorama: scaled expression data (HVGs only) was used. The wrapper function scanpy.external.pp.scanorama_integrate in Scanpy was used to generate the latent representations with the dataset of origin being a batch confounder, followed by neighborhood construction (scanpy.pp.neighbors) using the derived latent space and UMAP generation (scanpy.tl.umap).
6) scVI: raw count expression (HVGs only) was used with the batch covariate being the dataset of origin. A scVI model was built using the function scvi.model.SCVI (tag-value pairs: n_layers = 2, n_latent = 30, gene_likelihood = "nb") and further trained to generate the latent representations, followed by neighborhood construction (scanpy.pp.neighbors) using the derived latent space and UMAP generation (scanpy.tl.umap).
7) scANVI: raw count expression (HVGs only) was used with the batch covariate being the dataset of origin. The function scvi.model.SCVI was first applied as in scVI, and a scANVI model was next built using the function scvi.model.SCANVI.from_scvi_model (the biological key is set as the cell annotations derived from the CellHint harmonization procedure; both high- and low-hierarchy annotations were used). This model was trained with 20 epochs to generate the latent representations, followed by neighborhood construction (scanpy.pp.neighbors) using the derived latent space and UMAP generation (scanpy.tl.umap).
8) scGen: normalized expression (HVGs only) was used with the batch covariate being the dataset of origin and the biological covariate (high- or low-hierarchy annotations) being the cell annotations derived from the CellHint harmonization procedure. The function scgen.SCGEN in scGen was used to build the model, which after training, resulted in the latent representations, followed by neighborhood construction (scanpy.pp.neighbors) using the derived latent space and UMAP generation (scanpy.tl.umap).

### Benchmarking analysis of CellHint

We used the metrics summarized in scIB^74^ (version 1.1.1) to quantify the degree of biological conservation (average silhouette width [ASW], isolated label score ASW, and isolated label score F1) and batch removal (batch ASW, graph connectivity, and kBET score). Local inverse Simpson’s index metrics were not selected, and other metrics requiring a cluster key were also not used here. Due to an elongated runtime, only 50,000 cells were randomly sampled for assessing the six metrics. Considering that not all tools produce batch-corrected low-dimensional latent representations, we used the ultimate UMAP embeddings (see the previous section), together with the neighborhood graph constructed from the UMAP, to directly assess biological conservation and batch removal at the level of UMAP. Cells from the five immune datasets were next downsampled to 10,000, 50,000, 100,000, 200,000, 300,000 or all cells (417,866). The integration procedures as detailed in the Methods section “*Integration of datasets with different tools*” were followed for each cell subset. For scVI, scANVI and scGen, both GPU (NVIDIA Tesla T4) and CPU (Intel(R) Xeon(R) Gold 6226R CPU @ 2.90GHz) were utilized.

### Generation of multiple organ atlases

A total of 38 scRNA-seq and snRNA-seq datasets were collected. Detailed information of these datasets can be found in **Table S2**. Each tissue or organ consisted of four datasets. Because some datasets contained multiple organs, these datasets were repeatedly used in different tissue atlases. For each dataset, raw count expression data was downloaded, accompanied by the meta-information of original annotations, experimental assays, donor identities, ages, genders, and suspension types. These metadata were next unified across datasets.

Using the procedure in the Methods section “*Cell type harmonization*”, we performed automatic cell type harmonization for cells from each organ that comprised four datasets. After harmonization, we injected the cell reannotation information into the combined dataset for supervised data integration. For data integration, we used the annotation-aware procedure as detailed in the Methods section “*Annotation-aware data integration*”, with the batch key being the dataset of origin (for organs of heart, lung, and pancreas) or the donor ID (for organs of blood, bone marrow, hippocampus, intestine, kidney, liver, lymph node, skeletal muscle and spleen) depending on how these batches manifested in the organ studied. We next renamed the harmonized cell types into biologically meaningful labels, followed by further manual curations of cell annotations including missed cell subtypes. These cell types were also linked with the cell type labels from the Cell Ontology^75^ for future community-wide query.

Machine learning-based CellTypist models were next built for each tissue and each dataset from the given tissue. The function celltypist.train in CellTypist was used to train logistic regression-based classifiers, with the manual annotation being the label key and feature selection turned on (tag-value pairs: feature_selection = True, n_top_genes = 300). Ultimately, five models were trained for each tissue (four dataset-specific models and one combined model), resulting in a total of 60 models that could be used for automatic cell type annotation for cells querying our database in different tissue contexts. The integrated datasets with harmonized and curated cell annotations in 12 organs, as well as the CellTypist models, are all available at https://www.celltypist.org/organs.

## QUANTIFICATION AND STATISTICAL ANALYSIS

### Benchmarking analysis

To enable a fair comparison between CellHint and other tools, we used the default parameters of most tools or followed the recommended procedures in the official tutorials. One exception was the integration method in Seurat v4, where all pairwise anchors will be searched by default, resulting in much longer runtime. For this method, we specified the reference to be Domínguez Conde et al.^8^ and mapped all other datasets onto it. In Figure 5D, runtime of some tools was shown to decrease when the numbers of cells were increased, which was caused by the changes in data structure for convergence or by the dynamic setting of the maximum numbers of iterations according to data sizes. Metrics of biological conservation and batch removal were assessed after cell downsampling due to runtime considerations.

### Comparison across species

Considering that harmonization of cell types across species is not stable (especially for non-model organisms without good gene annotations), all related analyses in this study were conducted based on one-to-one gene orthologs. Of note, even for comparison between two species, such as that between humans and macaques in Figure S8G, we still adopted these one-to-one orthologs among four species. Specifically, we located 15,721, 16,019, and 16,763 Ensembl-based orthologs between humans and macaques, pigs, and mice, respectively. After overlapping them with the available genes in the single-cell datasets, ultimately we retained 7,885 genes for all cross-species analyses.

## ADDITIONAL RESOURCES

CellHint documentation is available at https://cellhint.readthedocs.io/en/latest, and related organ resource is at https://www.celltypist.org/organs.

## Supplemental information

Table S1. Detailed steps for cell type harmonization and integration using the CellHint algorithm, related to Figure 1

Table S2. List of organs and datasets used for constructing the multi-organ single-cell resource, related to Figure 7

## Notes

### Summary of Updates

Figure 6 added; new analyses relating to another human tissue; Software name change; Supplemental files reorganized

